# The Identification of Key Genes and Biological Pathways in Polycystic Ovary Syndrome and its Complications by Next Generation Squancing (NGS) and Bioinformatics Analysis

**DOI:** 10.1101/2022.12.29.522191

**Authors:** Basavaraj Vastrad, Chanabasayya Vastrad

## Abstract

Polycystic ovary syndrome (PCOS) is is associated with infertility, obesity, insulin resistance, hyperinsulinemia, type 2 diabetes mellitus, hypertension, cardiovascular problems, neurological and psychological problems and cancer. The specific mechanism of PCOS and its complications remains unclear. The aim of this study was to apply a bioinformatics approach to reveal related pathways or genes involved in the development of PCOS and its complications. The next generation squancing (NGS) datset GSE199225 was downloaded from the gene expression omnibus (GEO) database. Differentially expressed gene (DEG) analysis was performed using DESeq2. The g:Profiler was utilized to analyze the functional enrichment, gene ontology (GO) and REACTOME pathway of the differentially expressed genes. A protein-protein interaction (PPI) network was constructed and module analysis was performed using HiPPIE and cytoscape. The miRNA-hub gene regulatory network and TF-hub gene regulatory network were also constructed for research. The expression of the hub genes was validated using receiver operating characteristic (ROC) curve analysis. We have identified 957 DEGs in total, including 478 up regulated genes and 479 down regulated gene. GO and REACTOME illustrated that DEGs in PCOS were significantly enriched in regulation of molecular function, developmental process, interferon signaling and platelet activation, signaling and aggregation. Finally, through analyzing the PPI network, modules, miRNA-hub gene regulatory network and TF-hub gene regulatory network, we screened hub genes HSPA5, PLK1, RIN3, DBN1, CCDC85B, DISC1, AR, MTUS2, LYN and TCF4 by the Cytoscape software. This study uses a series of bioinformatics technologies to obtain hug genes and key pathways related to PCOS and its complications. These analysis results provide us with novel ideas for finding biomarkers and treatment methods for PCOS and its complications.

## Introduction

Polycystic ovary syndrome (PCOS) is common reproductive endocrine disorders in women, characterized by hirsutism, anovulation, and polycystic ovaries [1]. It is estimated that the incidence of PCOS found in 5-20% of the female population [2]. PCOS is leading cause of infertility and adverse pregnancy outcomes in women [3]. The cause of the onset of it includes genetic and environmental factors [4]. Furthermore, PCOS might cause complications such as obesity [5], insulin resistance [6], hyperinsulinemia [7], type 2 diabetes mellitus [8], hypertension [9], cardiovascular problems [10], neurological and psychological problems [11] and cancer (breast, ovarian and endometrial) [12]. However, it is difficult to detect and diagnose the disease during the early period. Therefore, it is essential to explore potential diagnostic and prognostic biomarkers and therapeutic targets of early PCOS.

Previous investigations have shown that early diagnosis and treatment can help reduce the progressive infertility in PCOS patients. Presently, there are several possible PCOS biomarkers, such as sex hormone-binding globulin (SHBG) [13], vascular endothelial growth factor (VEGF) and HIF1 [14], FDX1 [15], CYP11A1, CYP17A1, and CYP19A1 [16], and KISS1 [17]. Extensive researched have shown signaling pathways in PCOS included PI3K/Akt signaling pathway [18], androgen signaling pathway [19], hippo signaling pathway [20], NF- κB signaling pathway [21] and insulin signaling pathway [22]. However, there are only a less screening biomarkers and therapeutic interventions that are of importance for the clinical treatment of PCOS. Therefore, the clarification of more unique PCOS biomarkers is crucially needed for precisely identifying patients and developing therapies.

With the development of next generation sequencing (NGS) technologies, bioinformatics analyses have been widely occupied to identify disease-specific biomarkers, explore significant epigenetic and genetic modifications, and reveal the molecular pathogenesis of PCOS [23]. Therefore, a large number of beneficial clues could be marked for new investigation on the base of NGS data. Furthermore, many bioinformatics investigation on PCOS have been produced in recent years [24], which proved that the integrated bioinformatics methods could help us to further investigation and better exploring the underlying molecular mechanisms.

We chosen NGS dataset GSE199225 [25] from Gene Expression Omnibus (GEO) (https://www.ncbi.nlm.nih.gov/geo/) [26] database. We then applied for DESeq2 package in R software to obtain the differentially expressed genes (DEGs) in the NGS dataset. Furthermore, Gene Ontology (GO) and REACTOME pathway enrichment analysis were used to determine the potential functions of the biomarker. The hub genes, miRNA and TFs were identified by protein-protein interaction (PPI) network, module analysis, miRNA-hub gene regulatory network and TF-hub gene regulatory network analysis. The predictive capability of the hub genes was analyzed by receiver operating characteristic (ROC) curve. The study probably revealed the molecular pathogenesis and potential therapeutic target of PCOS and its complications.

## Materials and Methods

### Next generation sequencing data source

NGS dataset GSE199225 [25] was acquired from the GEO database. The GSE199225 profile was composed of 30 PCOS samples and 30 normal control samples. GSE199225 dataset was based on GPL24676 Illumina NovaSeq 6000 (Homo sapiens).

### Identification of DEGs

Genes that were differentially expressed in PCOS samples were identified using the DESeq2 [27] package in R software, with DEGs being those genes with a P < 0.05, and a ∣logFC | > 0.375 for up regulated genes and ∣logFC | < -0.257 for down regulated genes. ggplot2 package in R software was used to construct volcano plot. log2-transformed mRNA expression data were arranged into heatmap using the gplot package in R software.

### GO and pathway enrichment analyses of DEGs

GO function (GO, http://www.geneontology.org) [28] and REACTOME (https://reactome.org/) [29] pathway enrichment analyses of the DEGs were performed using the g:Profiler (http://biit.cs.ut.ee/gprofiler/) [30]. GO classifies gene functions into biological processes (BP), cellular component (CC) and molecular function (MF). P value < 0.05 was defined statistically significant.

### Construction of the PPI network and module analysis

PPI network among all DEGs was established based on an online tool HiPPIE interactome database (http://cbdm-01.zdv.uni-mainz.de/~mschaefer/hippie/) [31]. Cytoscape (v 3.9.1) (http://www.cytoscape.org/) [32] was used to visualize the network, while the Network Analyzer plugin was used to rank genes within this network based upon their node degree [33], betweenness [34], stress [35] and closeness [36] score. In addition, the PEWCC1 [37] app in Cytoscape was used to check modules of the PPI network (degree cutoff = 2, max. Depth = 100, k- core = 2, and node score cutoff = 0.2).

### miRNA-hub gene regulatory network construction

To search for the targets of miRNAs, the miRNet database (https://www.mirnet.ca/) [38] online database was used. The genes identified in 14 databases (TarBase, miRTarBase, miRecords, miRanda (S mansoni only), miR2Disease, HMDD, PhenomiR, SM2miR, PharmacomiR, EpimiR, starBase, TransmiR, ADmiRE, and TAM 2) were considered as targets of miRNAs. The regulatory network between miRNAs and hub genes, which screened for predicted target genes, was further constructed to search for key molecules and axes in PCOS. The miRNA-hub gene regulatory network was visualized by Cytoscape 3.9.1 [32].

### TF-hub gene regulatory network construction

To search for the targets of TFs, the NetworkAnalyst database (https://www.networkanalyst.ca/) [39] online database was used. The genes identified in database (ChEa) were considered as targets of TFs. The regulatory network between TFs and hub genes, which screened for predicted target genes, was further constructed to search for key molecules and axes in PCOS. The TF-hub gene regulatory network was visualized by Cytoscape 3.9.1 [32].

### Receiver operating characteristic curve (ROC) analysis

ROC analysis was performed in different datasets to evaluate the diagnostic value of hub genes in PCOS. To evaluate the role of hub genes in the diagnosis of PCOS, ROC analysis was conducted with pROC package in R software [40]. The genes with area under curve (AUC) >0.8 were considered as indicated that the model had a good fitting effect.

## Results

### Identification of DEGs

The DEGs were screened by “DESeq2” package (p < 0.05 and ∣logFC | > 0.375 for up regulated genes and ∣logFC | < -0.257 for down regulated genes).The GSE199225 dataset contained 957 DEGs, including 478 up regulated genes and 479 down regulated genes and are listed in Table 1. DEGs identified by threshold were visualized by volcano plot (Fig. 1). DEGs ordered by adjusted p value in each sample were shown by heatmap (Fig. 2).

**Fig. 1.**
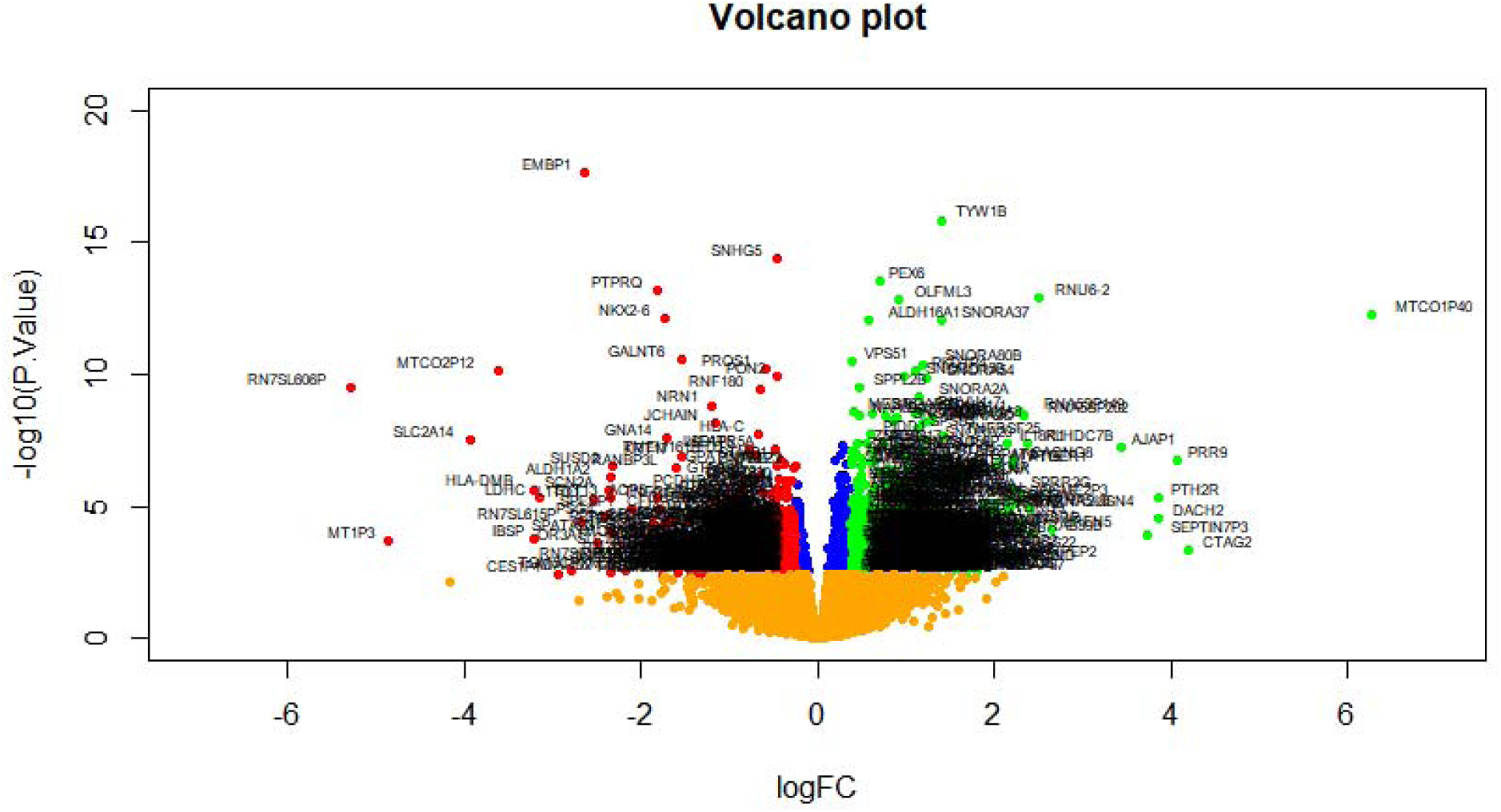
Volcano plot of differentially expressed genes. Genes with a significant change of more than two-fold were selected. Green dot represented up regulated significant genes and red dot represented down regulated significant genes.

**Fig. 2.**
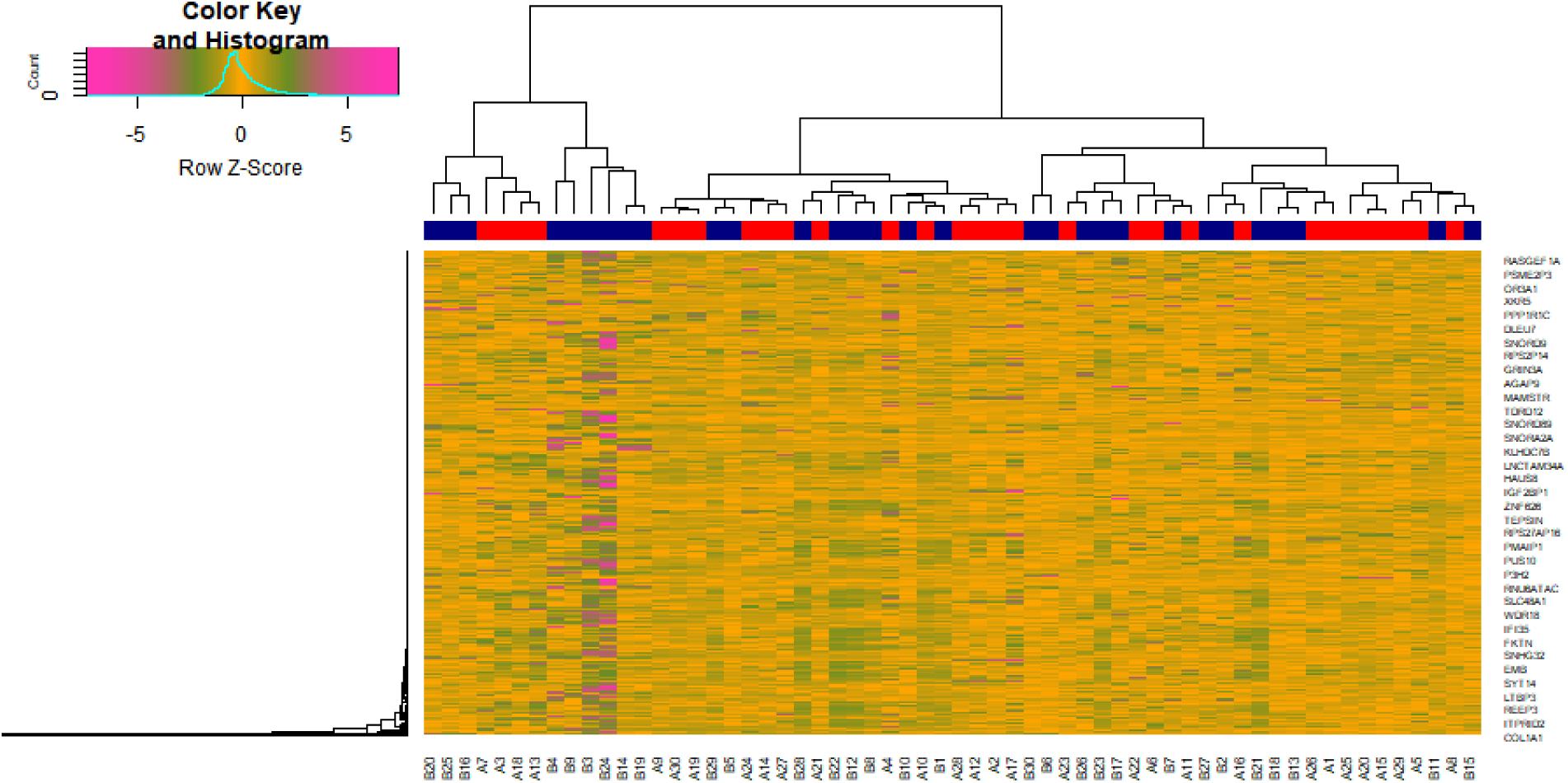
Heat map of differentially expressed genes. Legend on the top left indicate log fold change of genes. (A1 – A30 = normal control samples; B1 – B30 = PCOS samples)

**Table 1.**
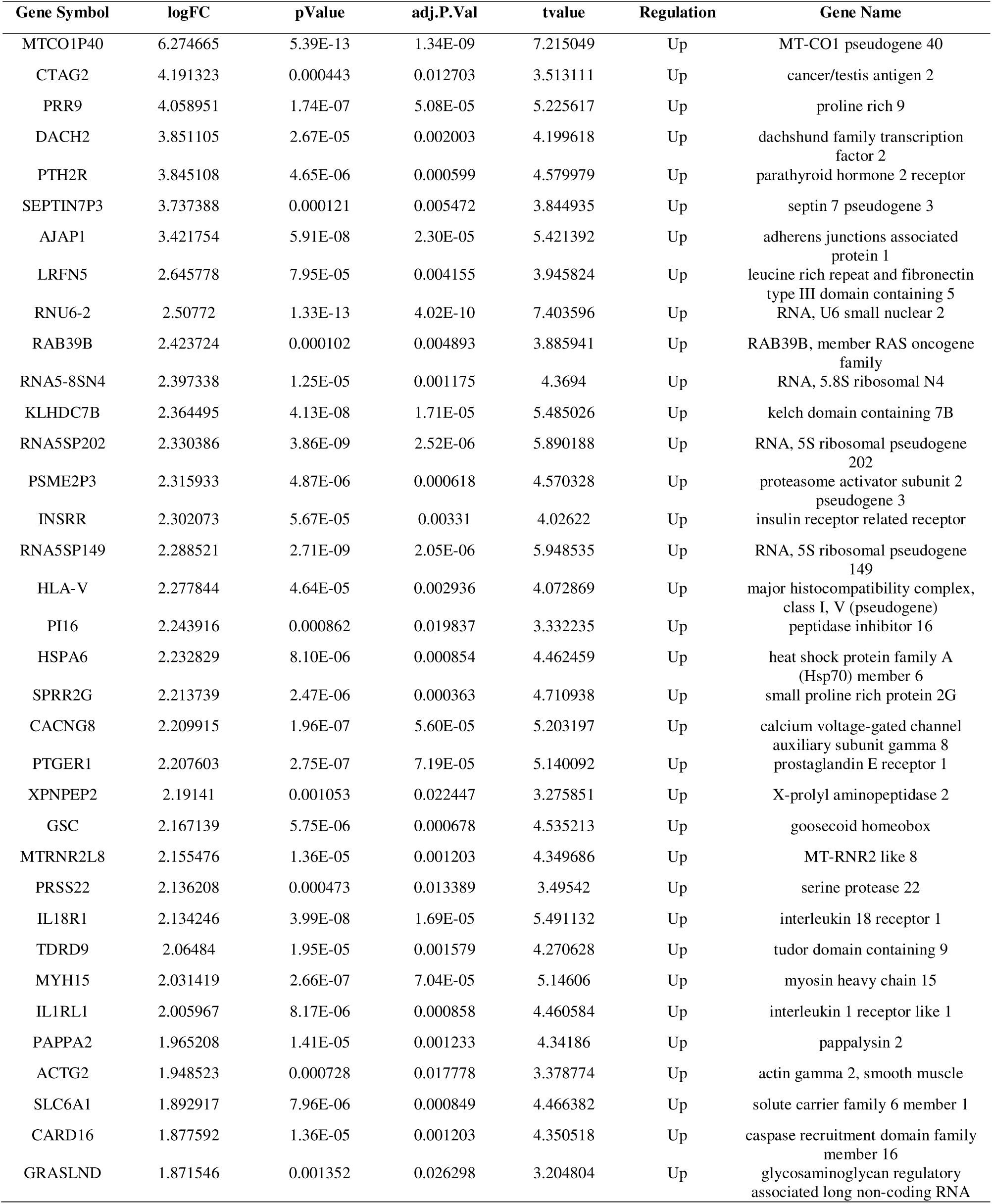
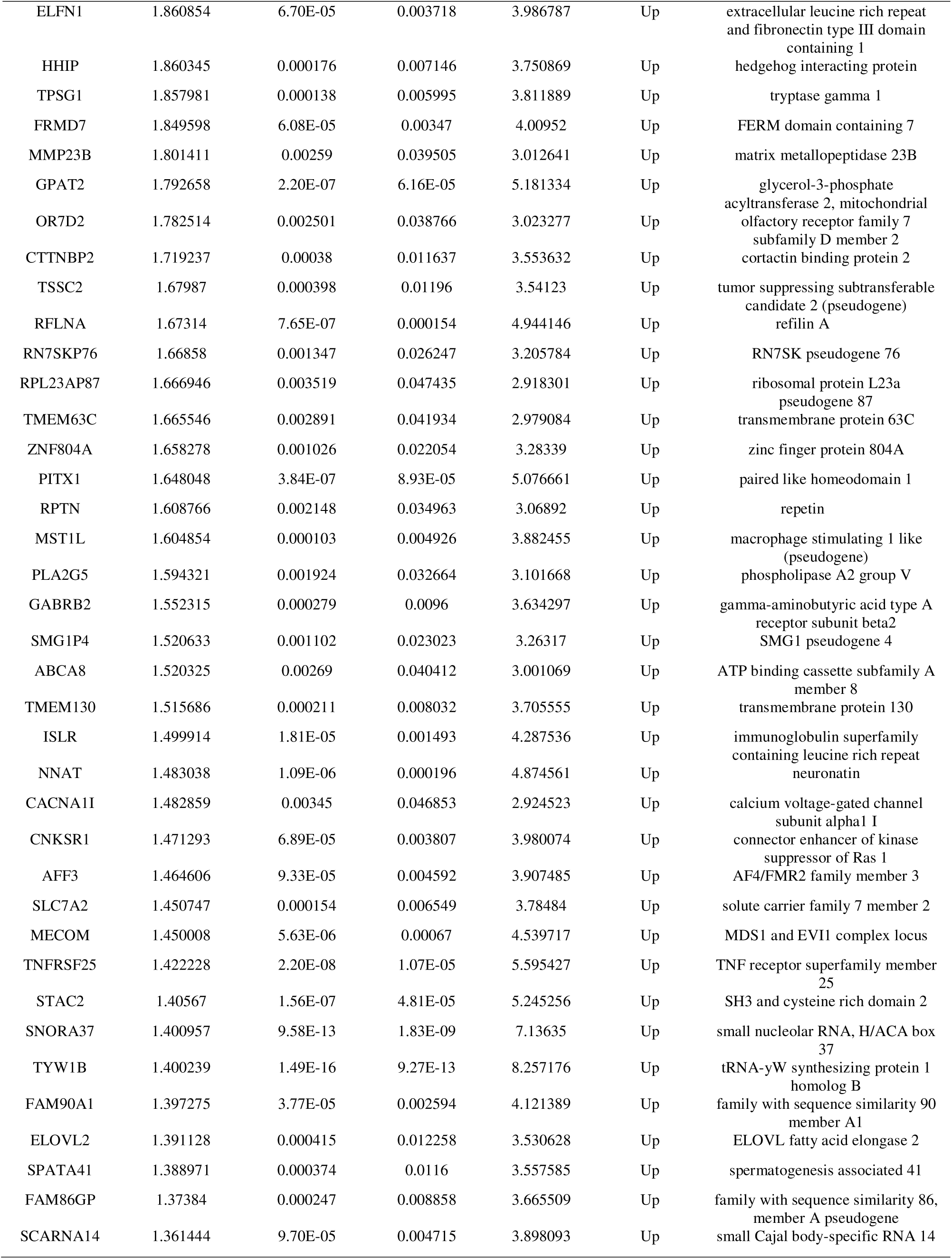
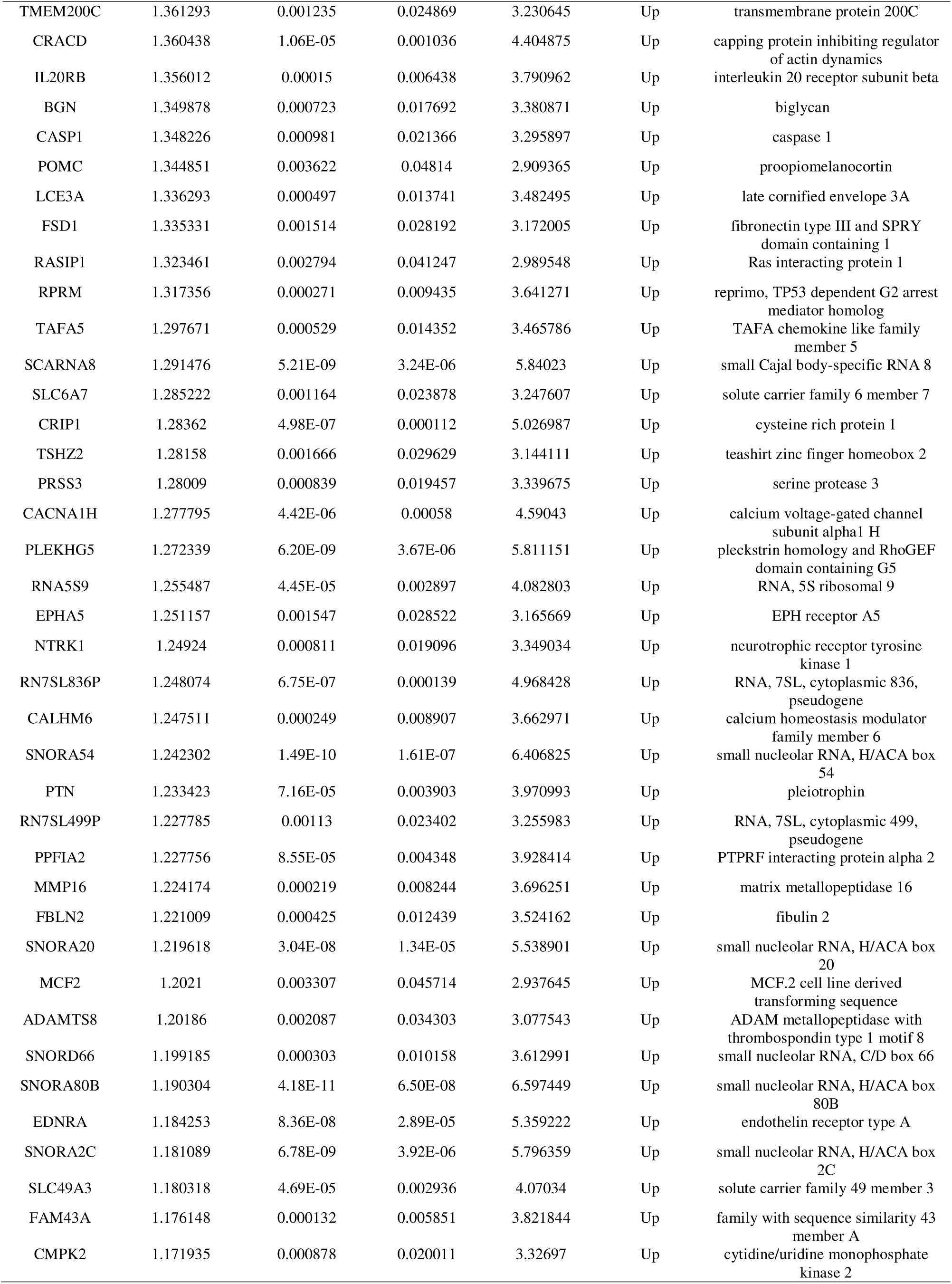
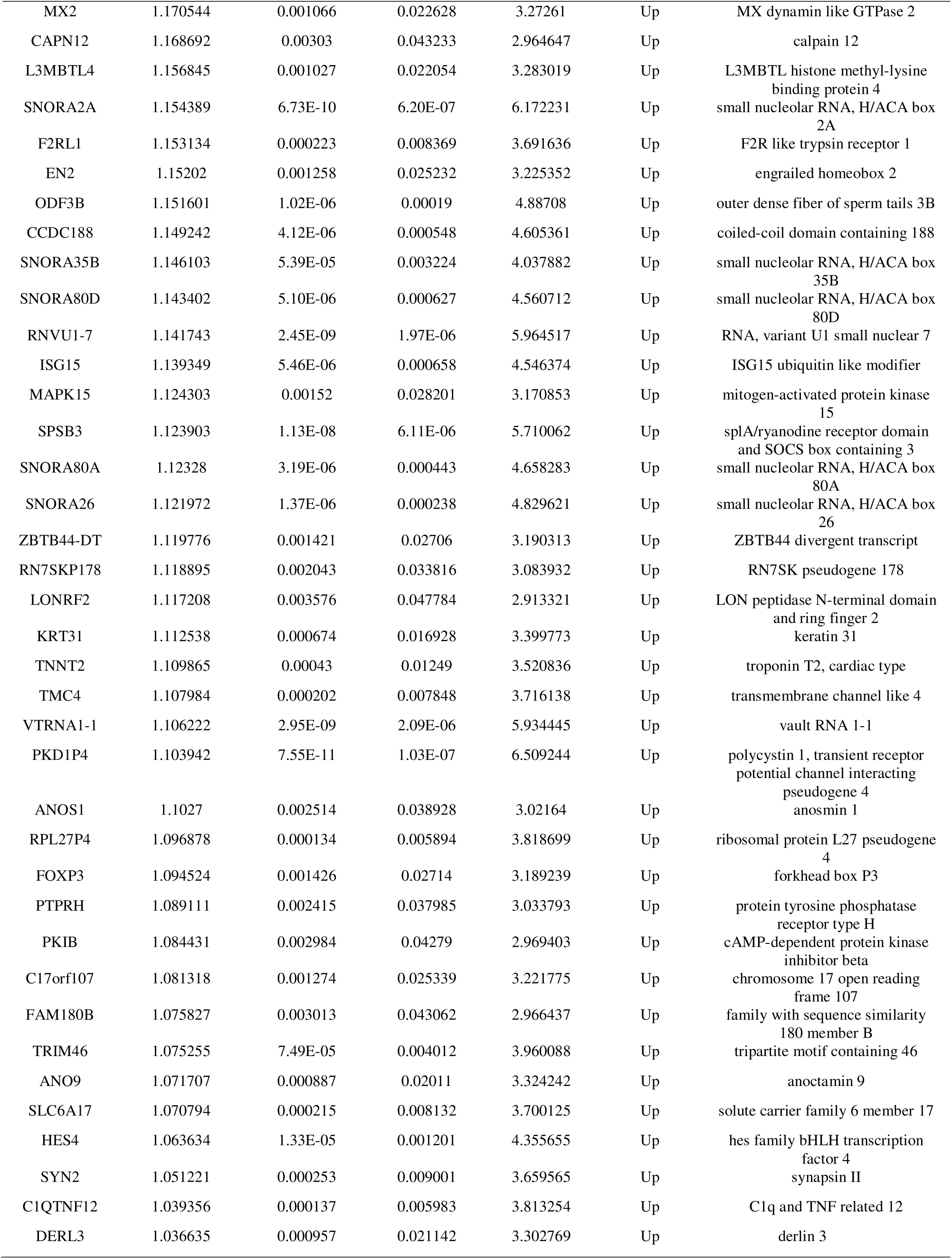
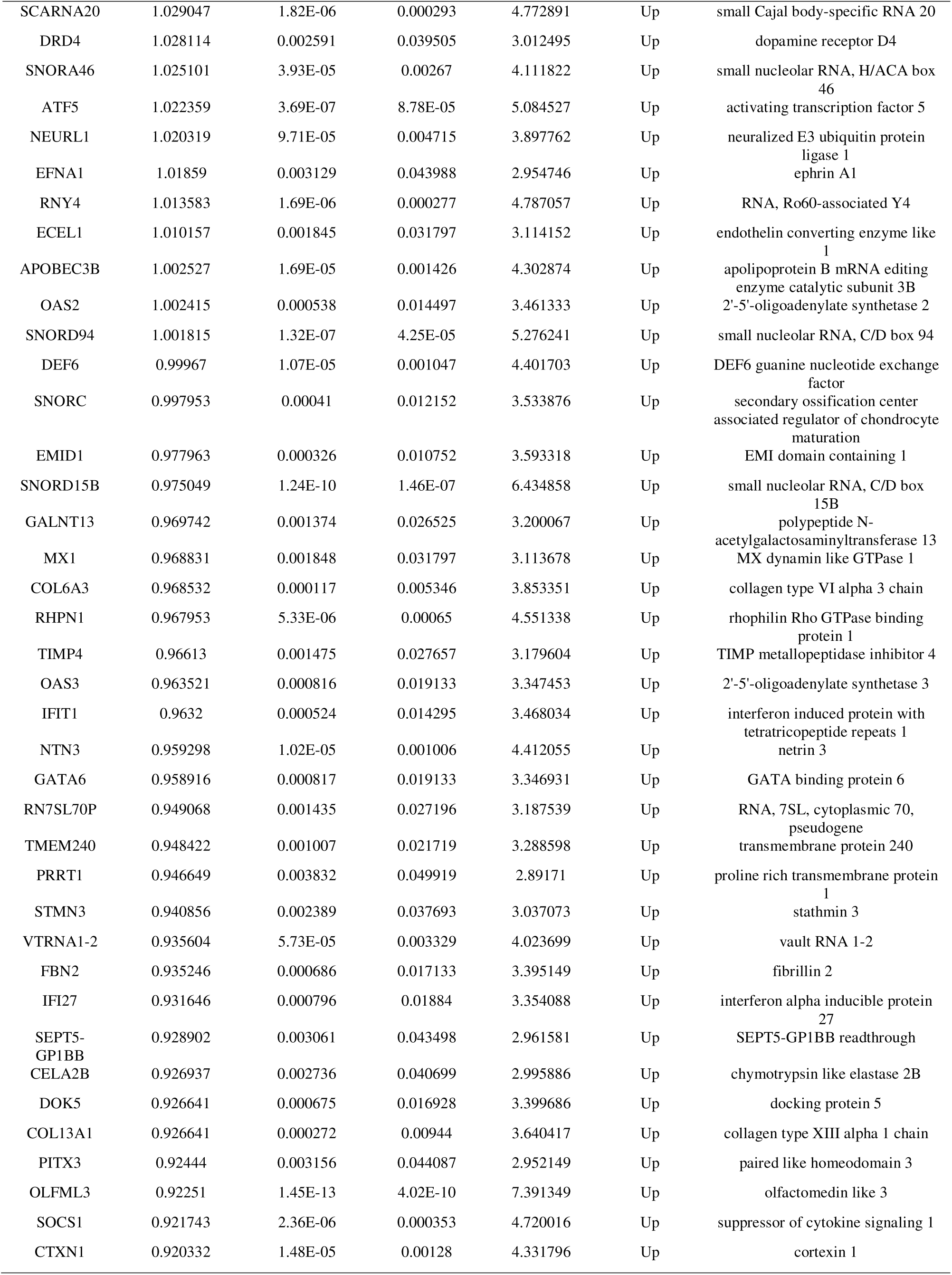
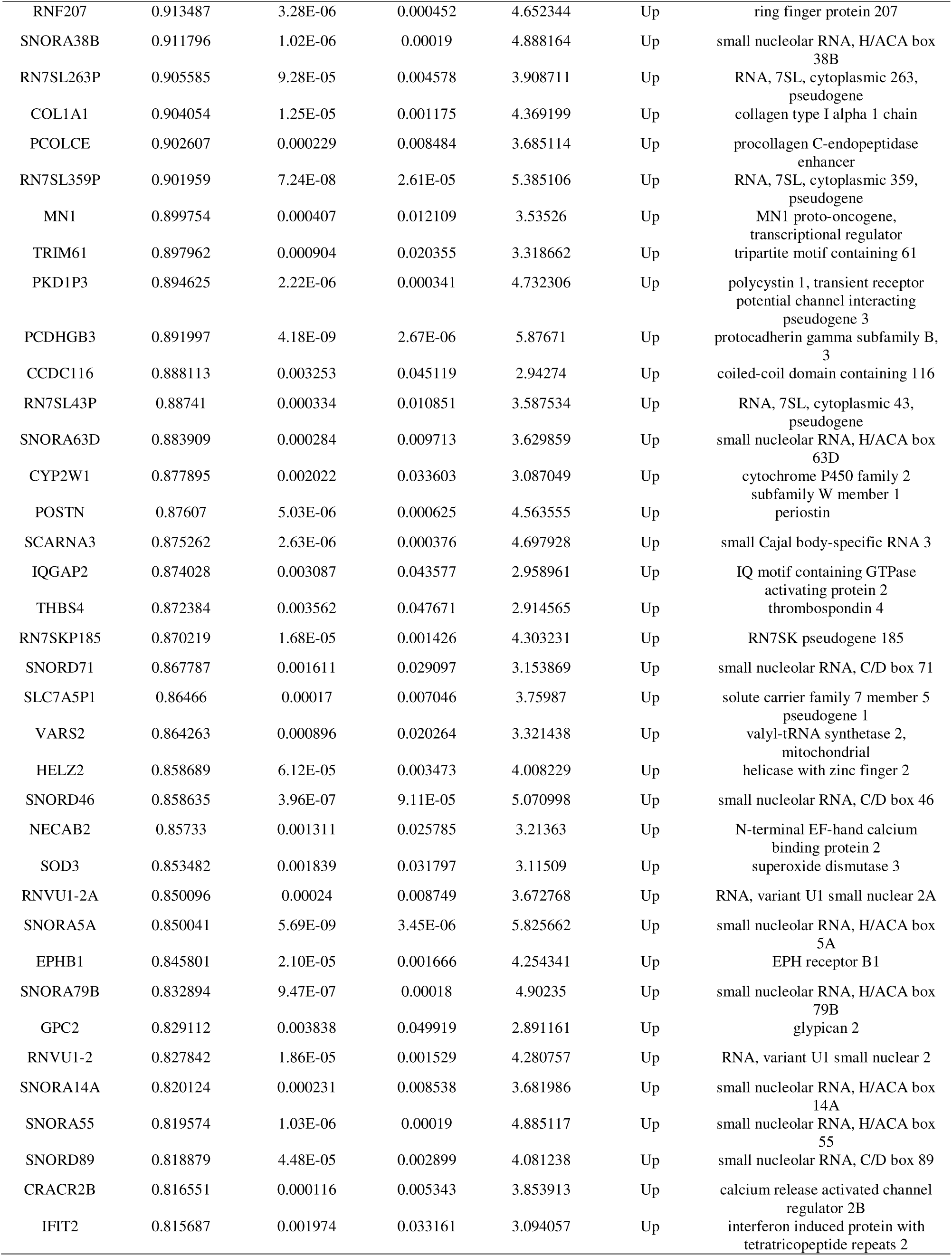
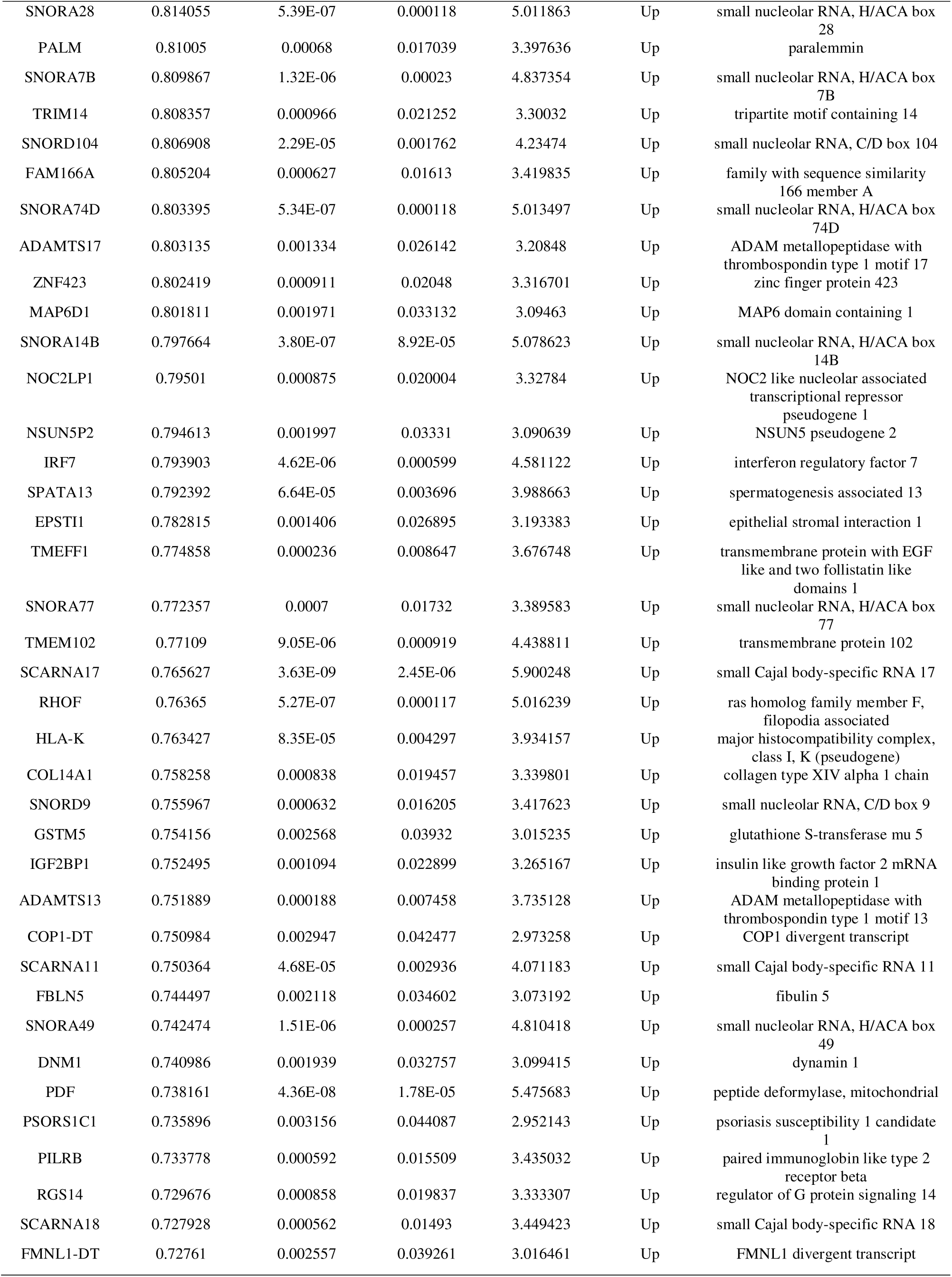
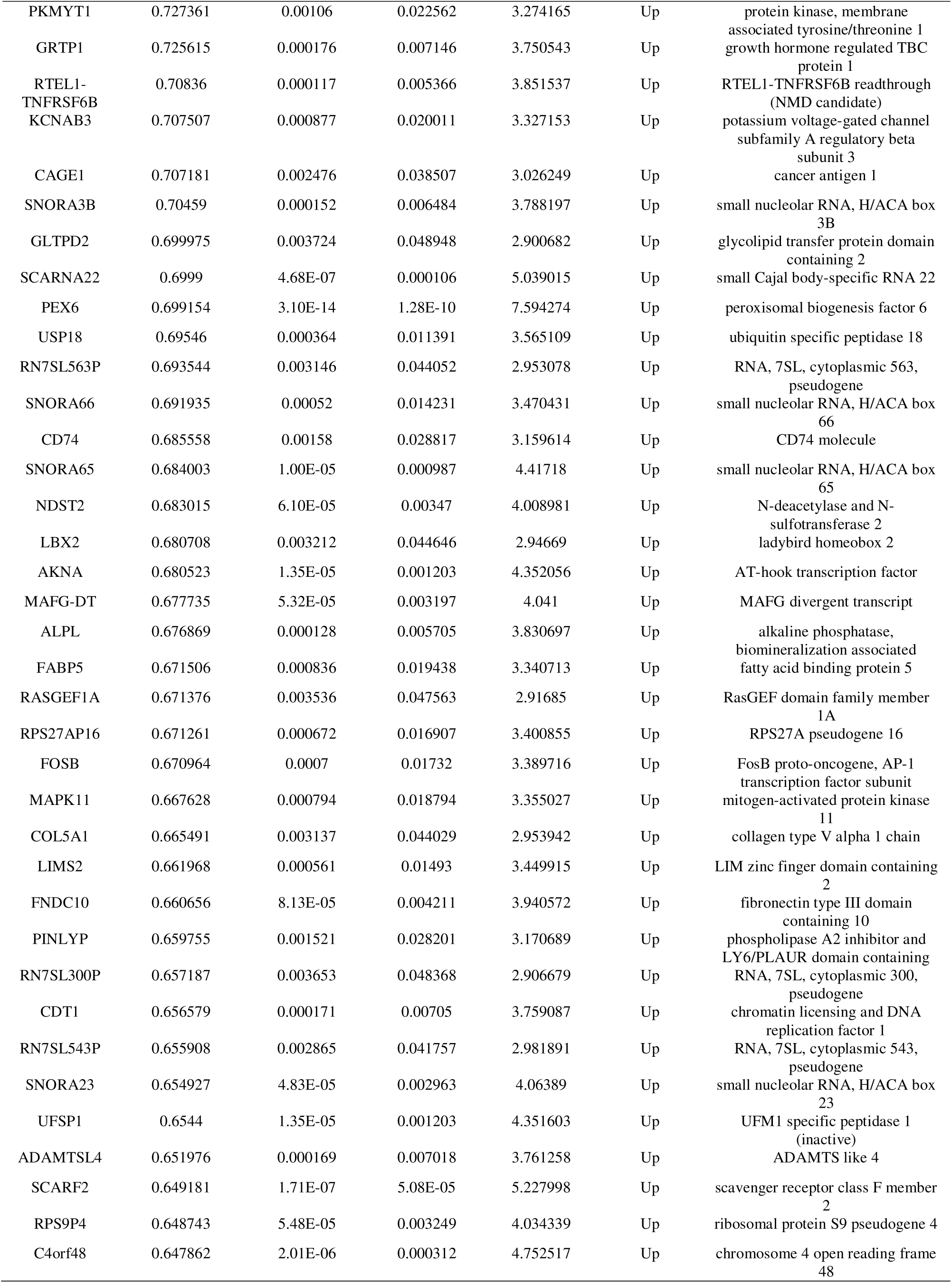
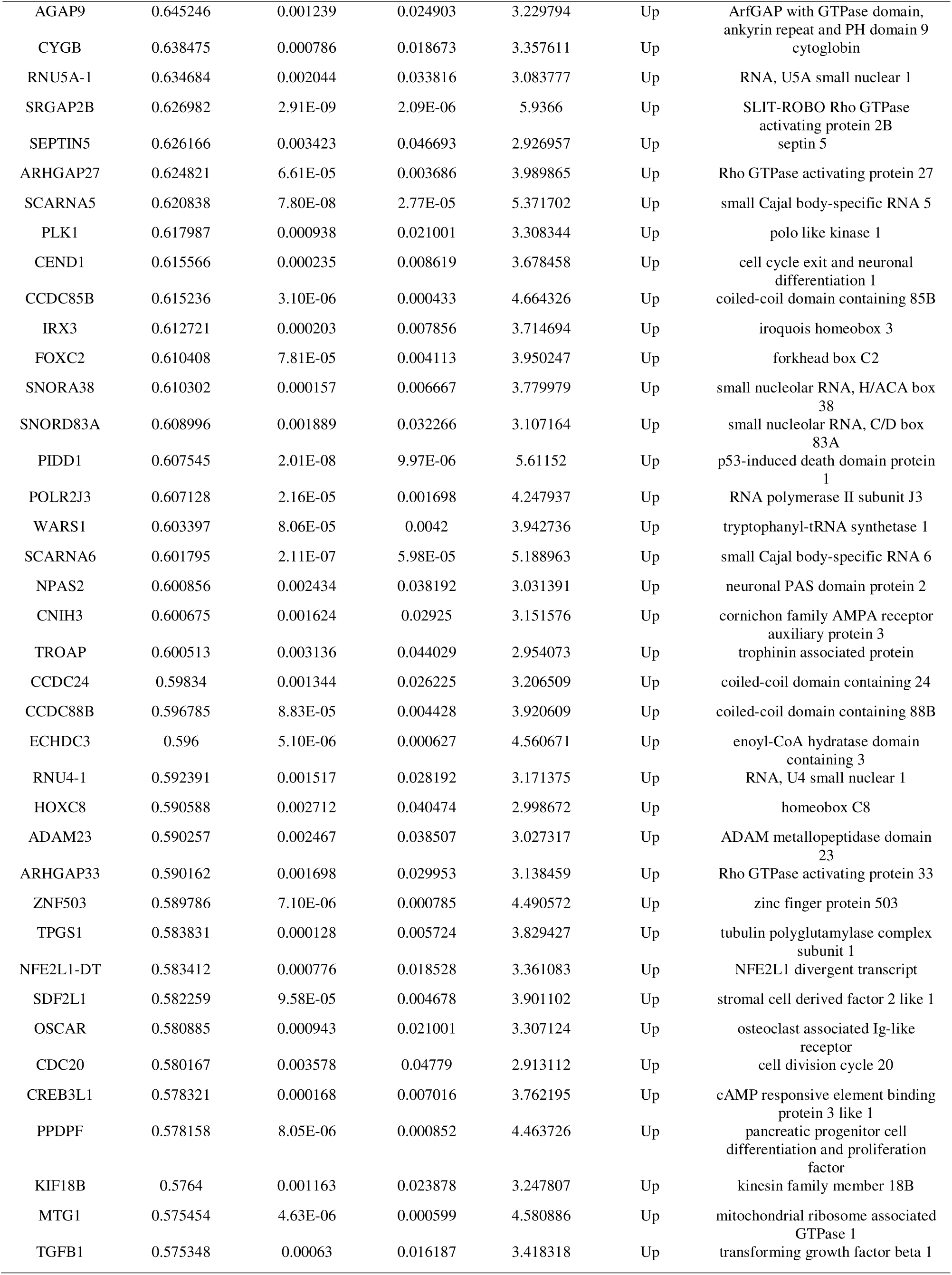
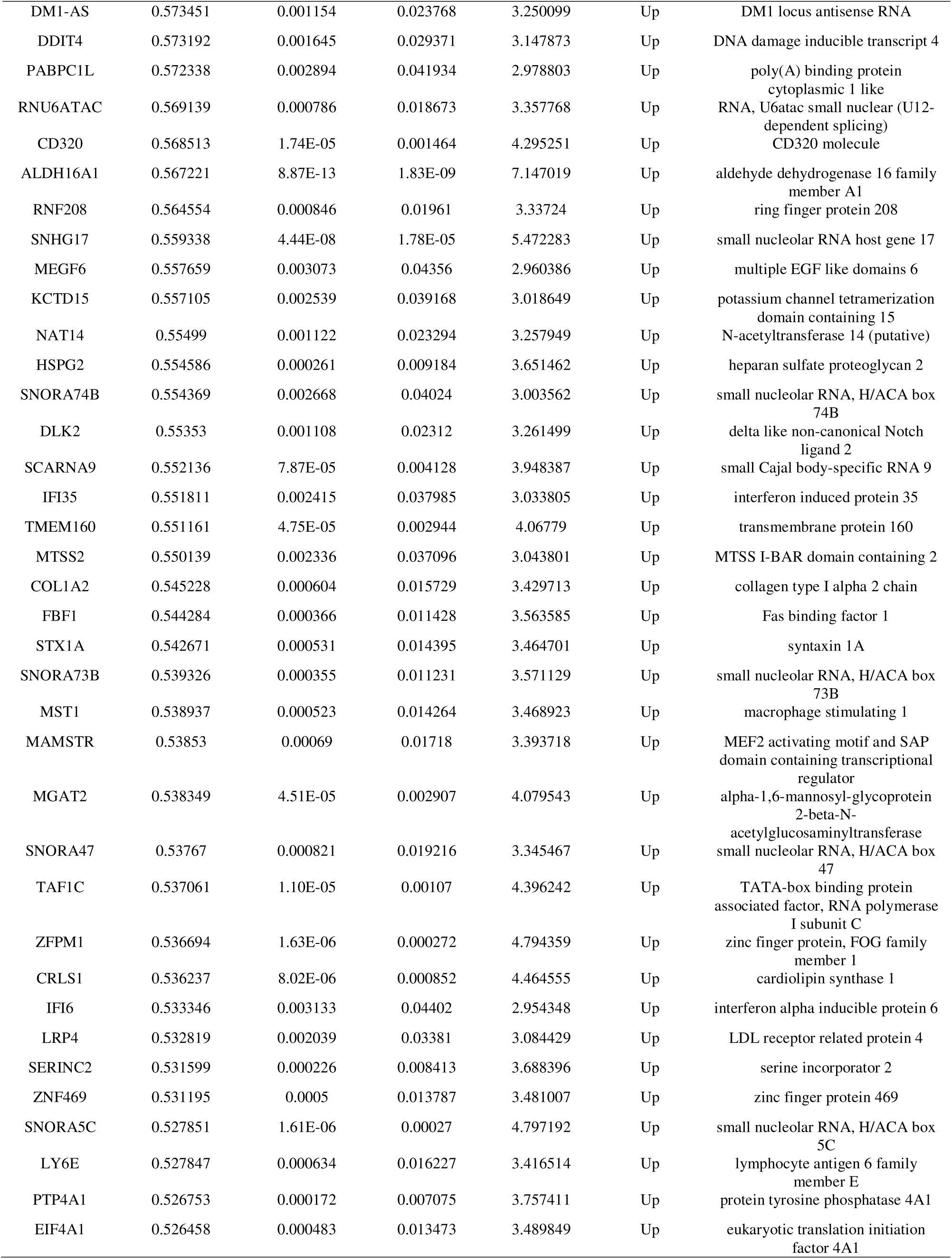
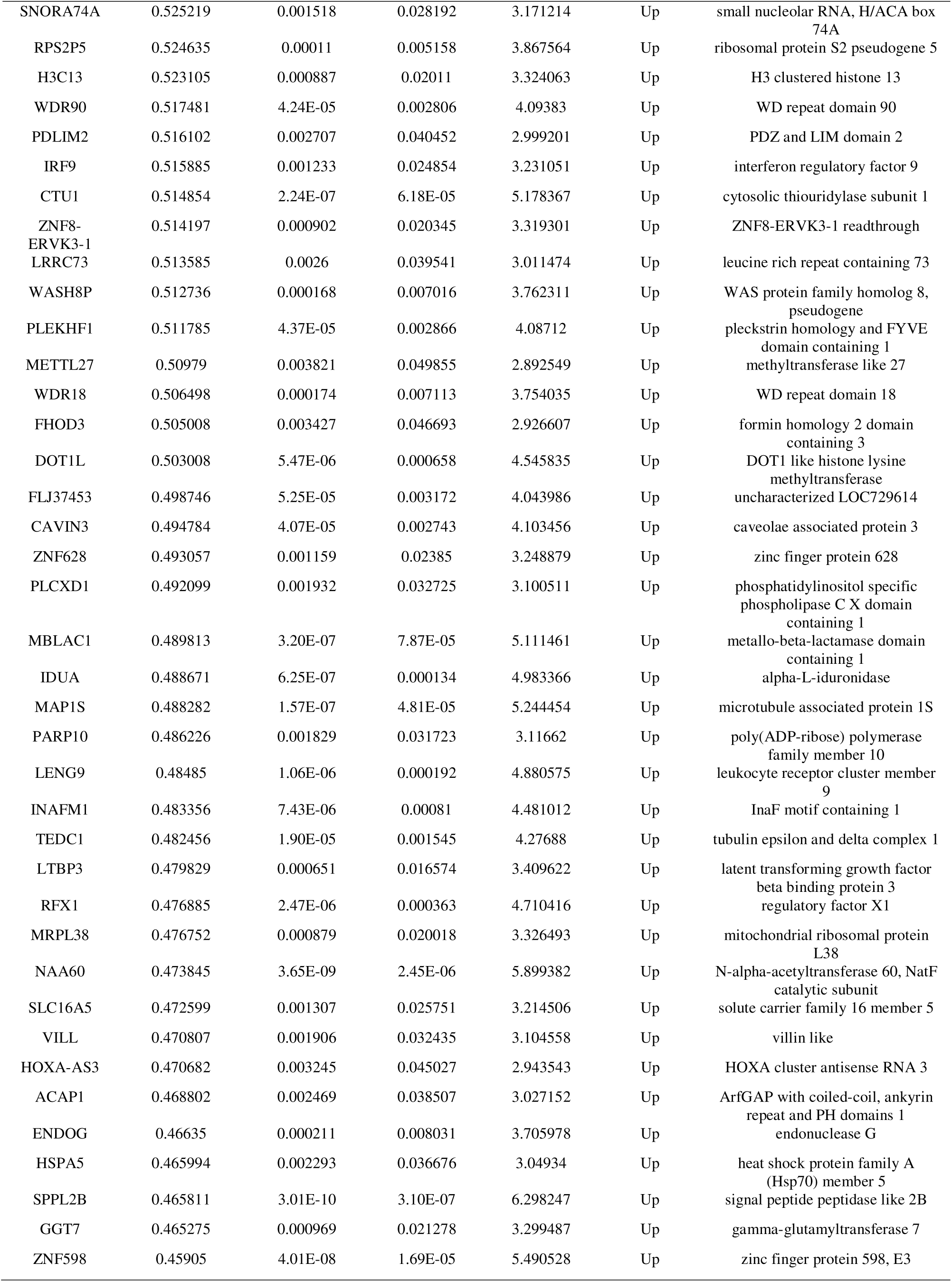
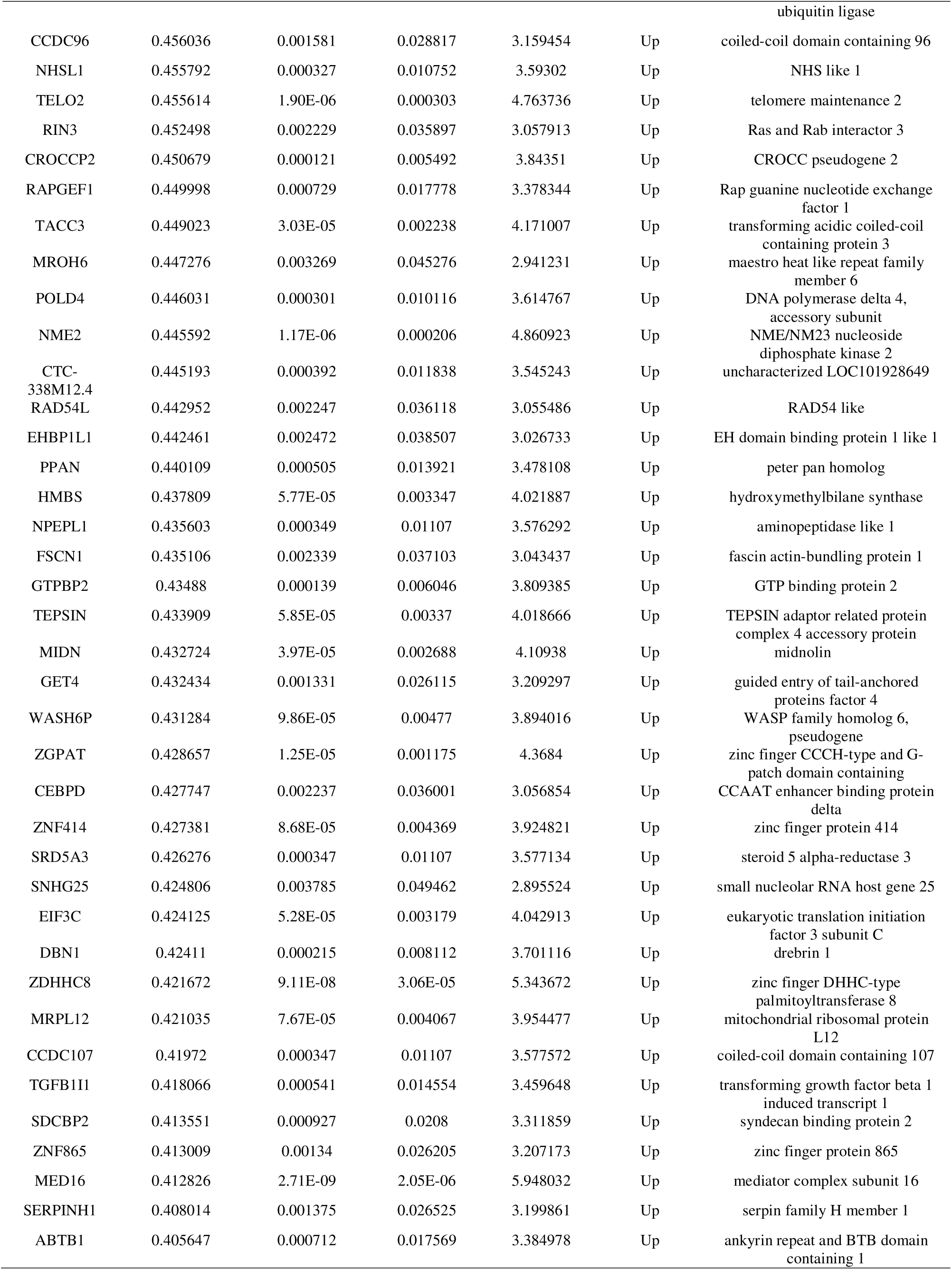
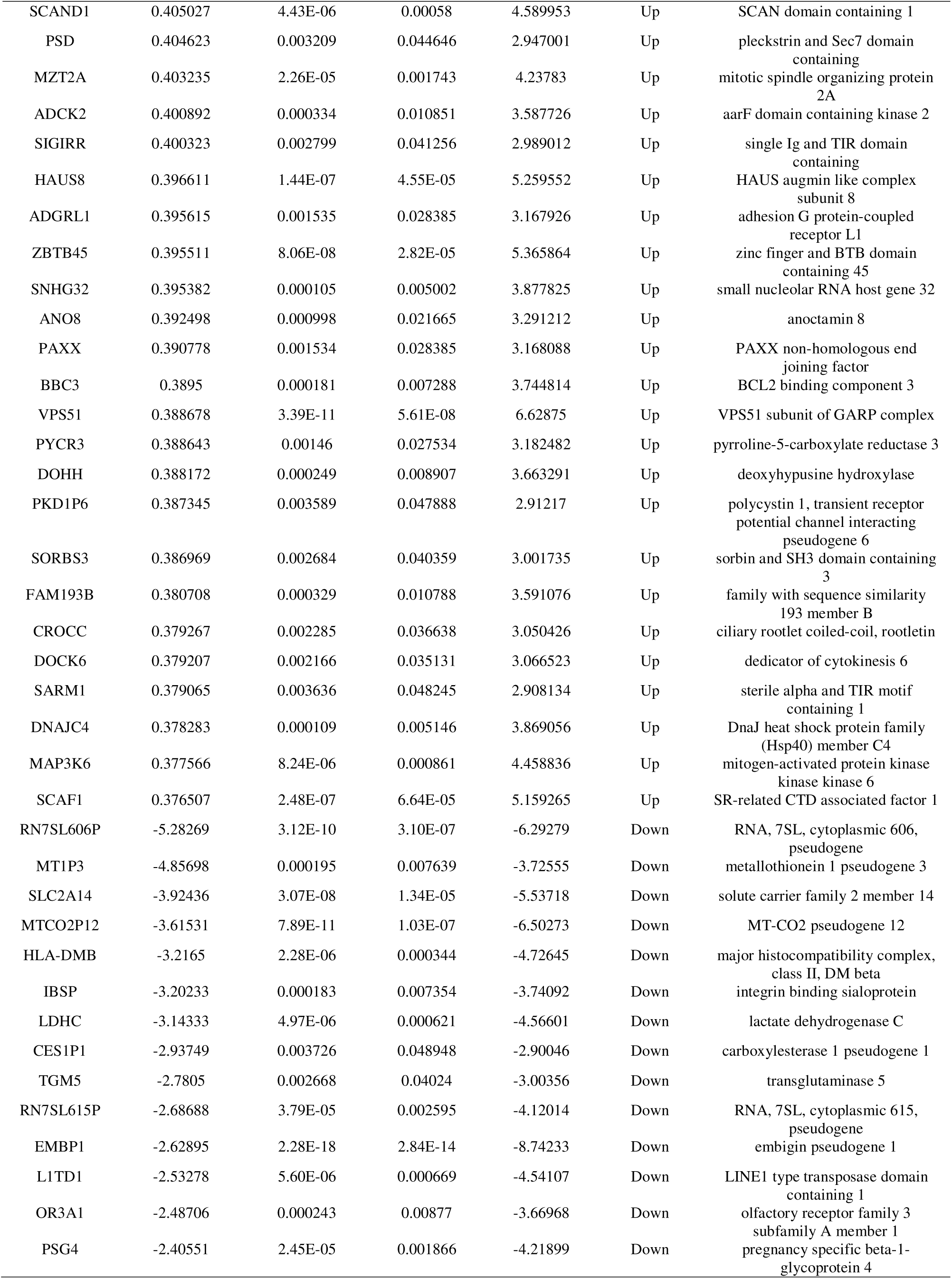
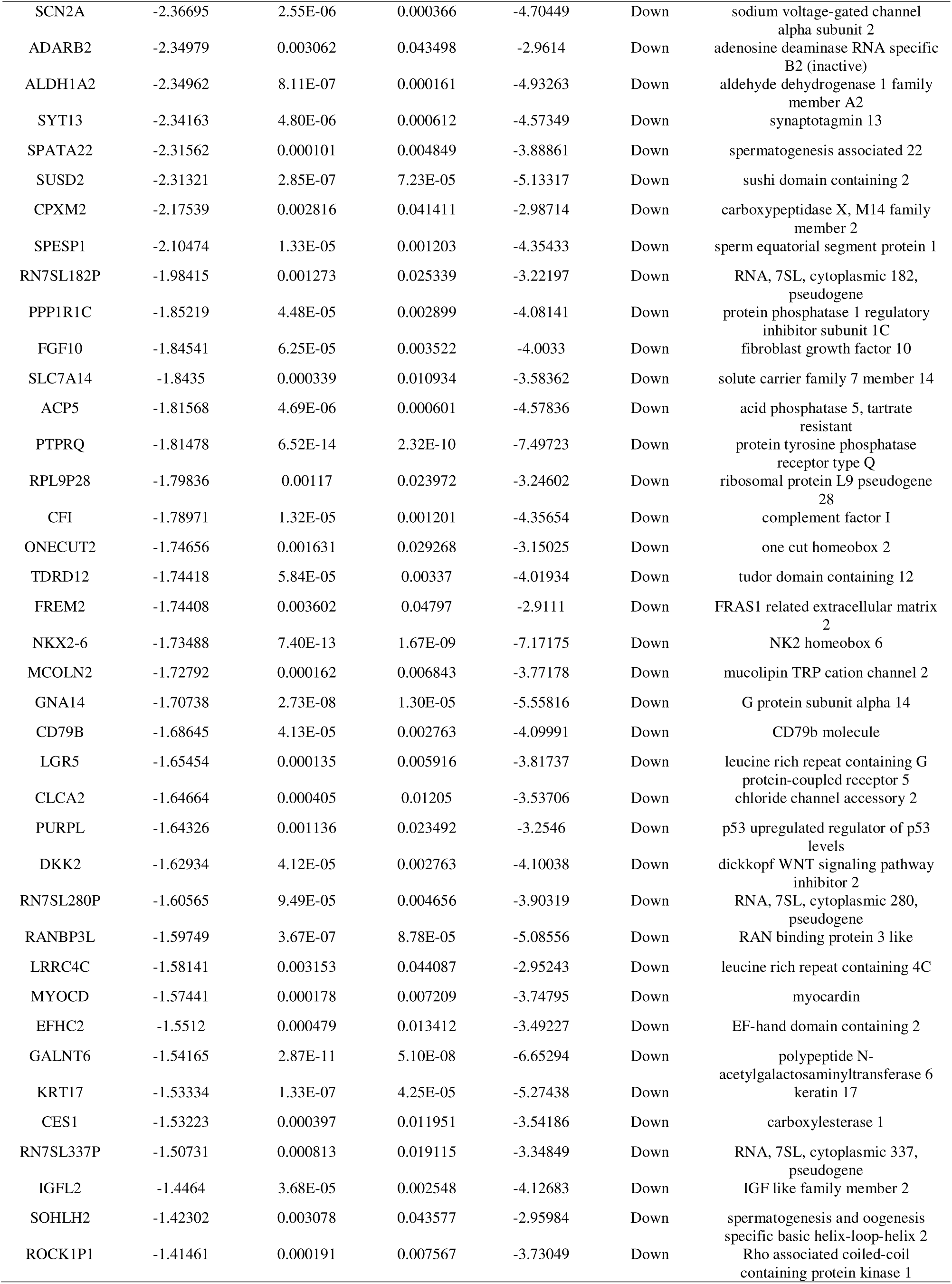
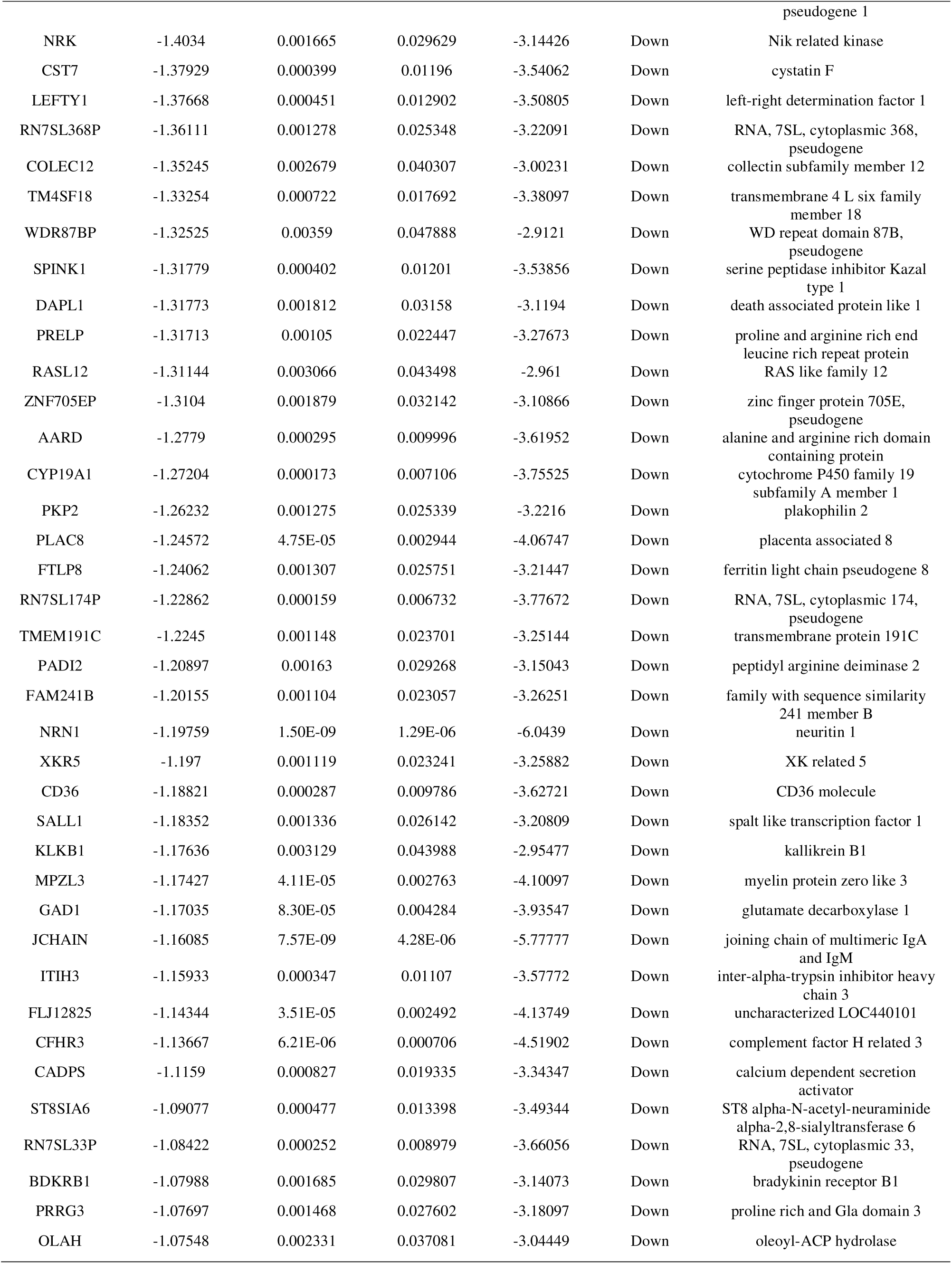
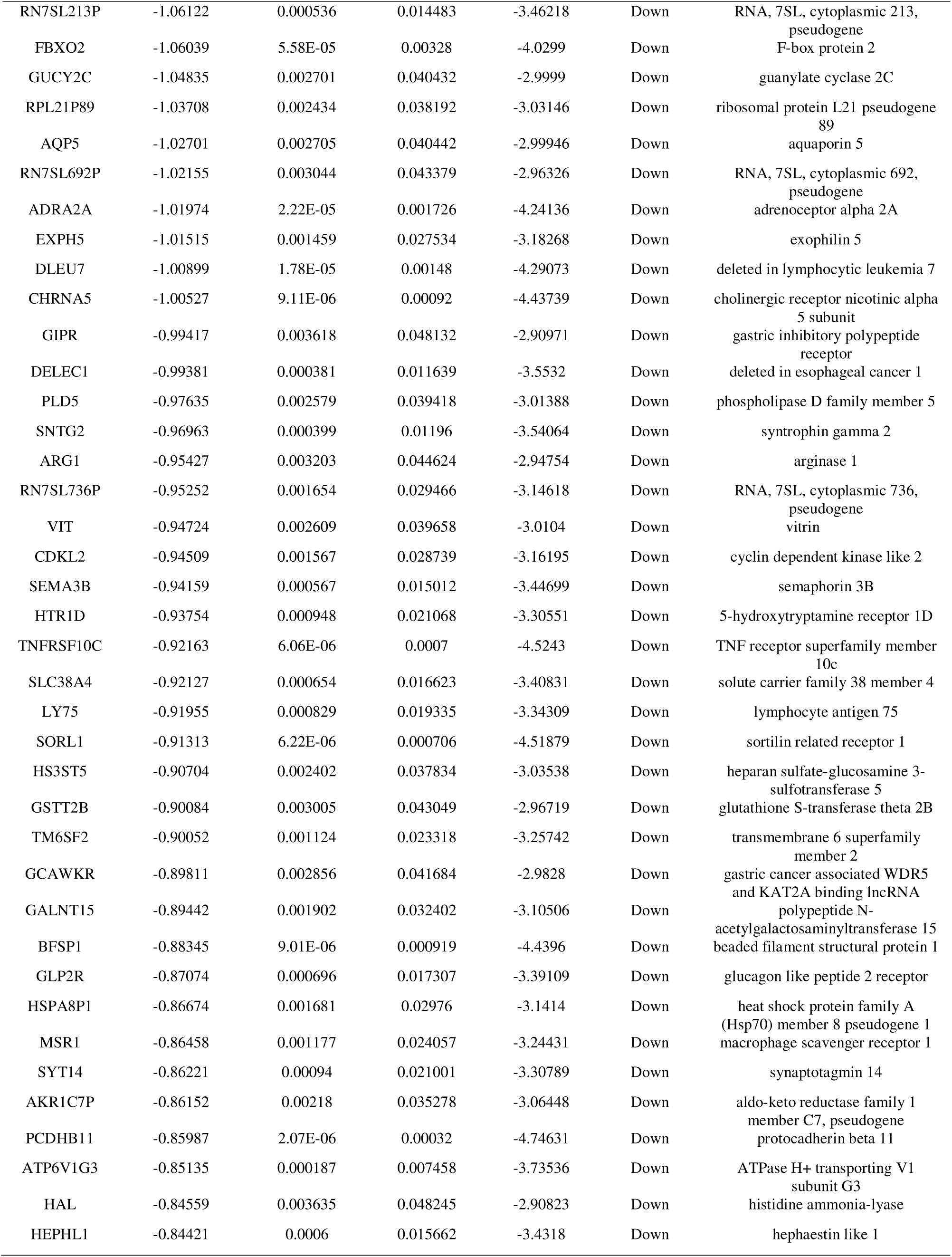
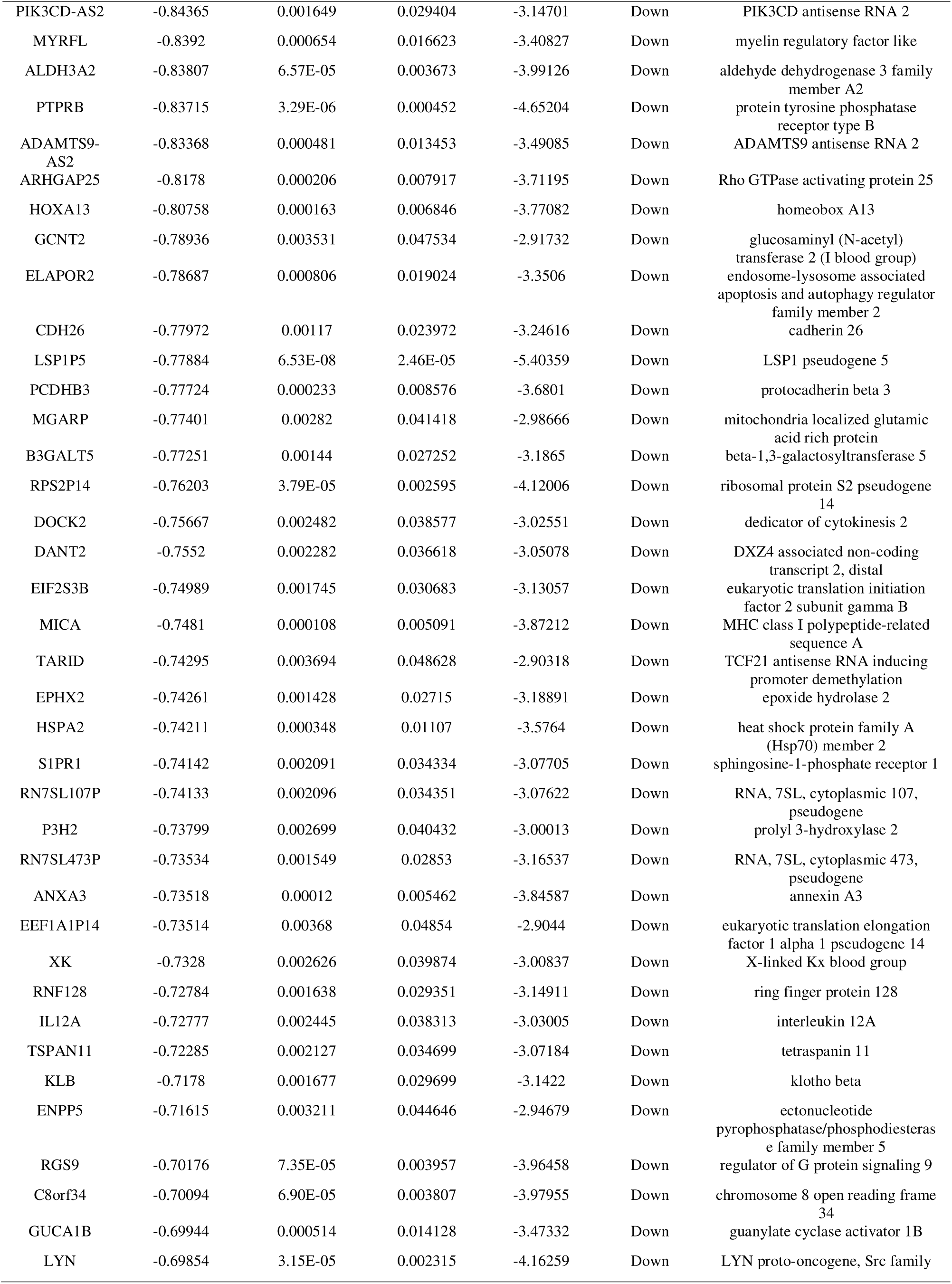
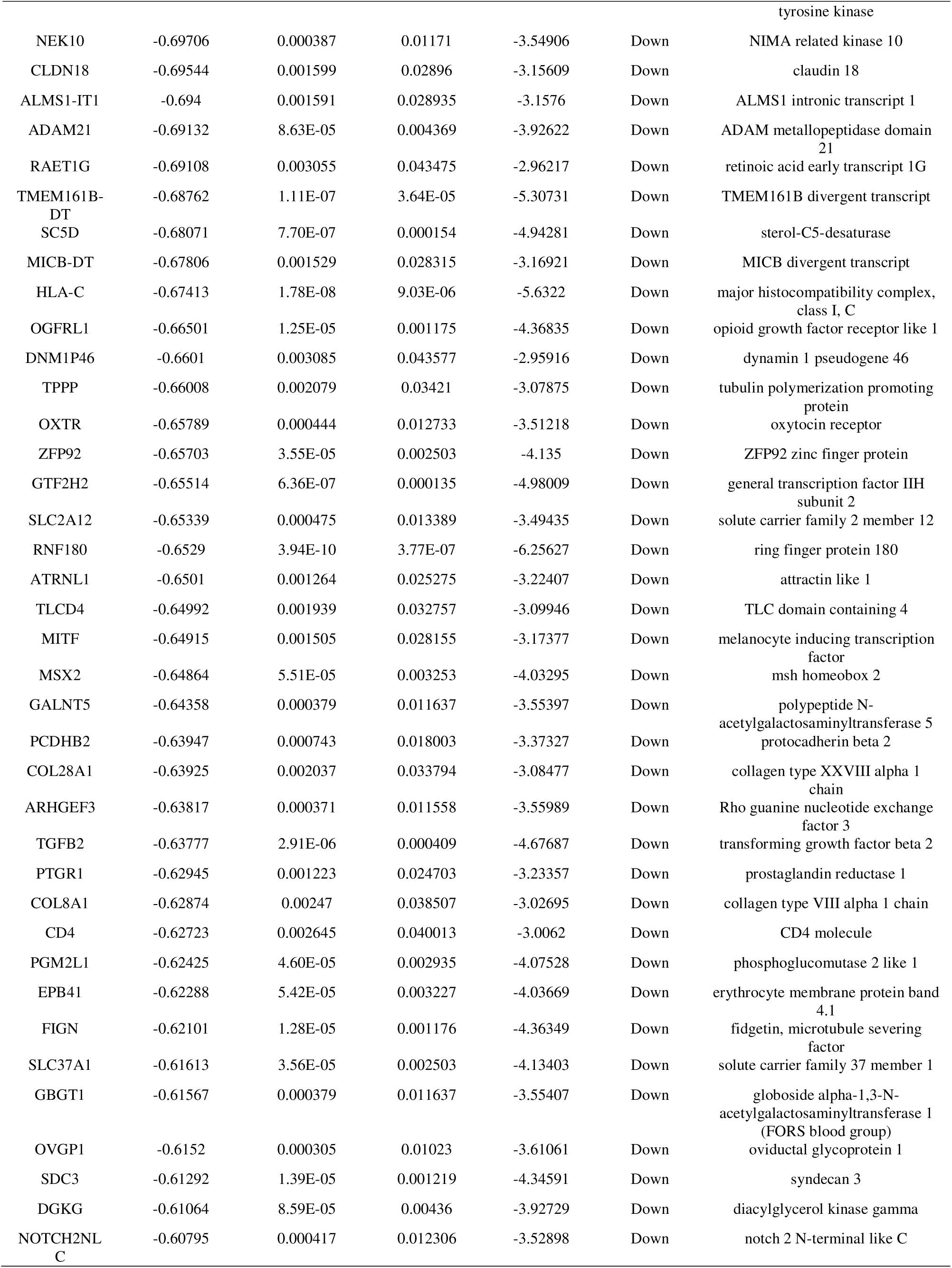
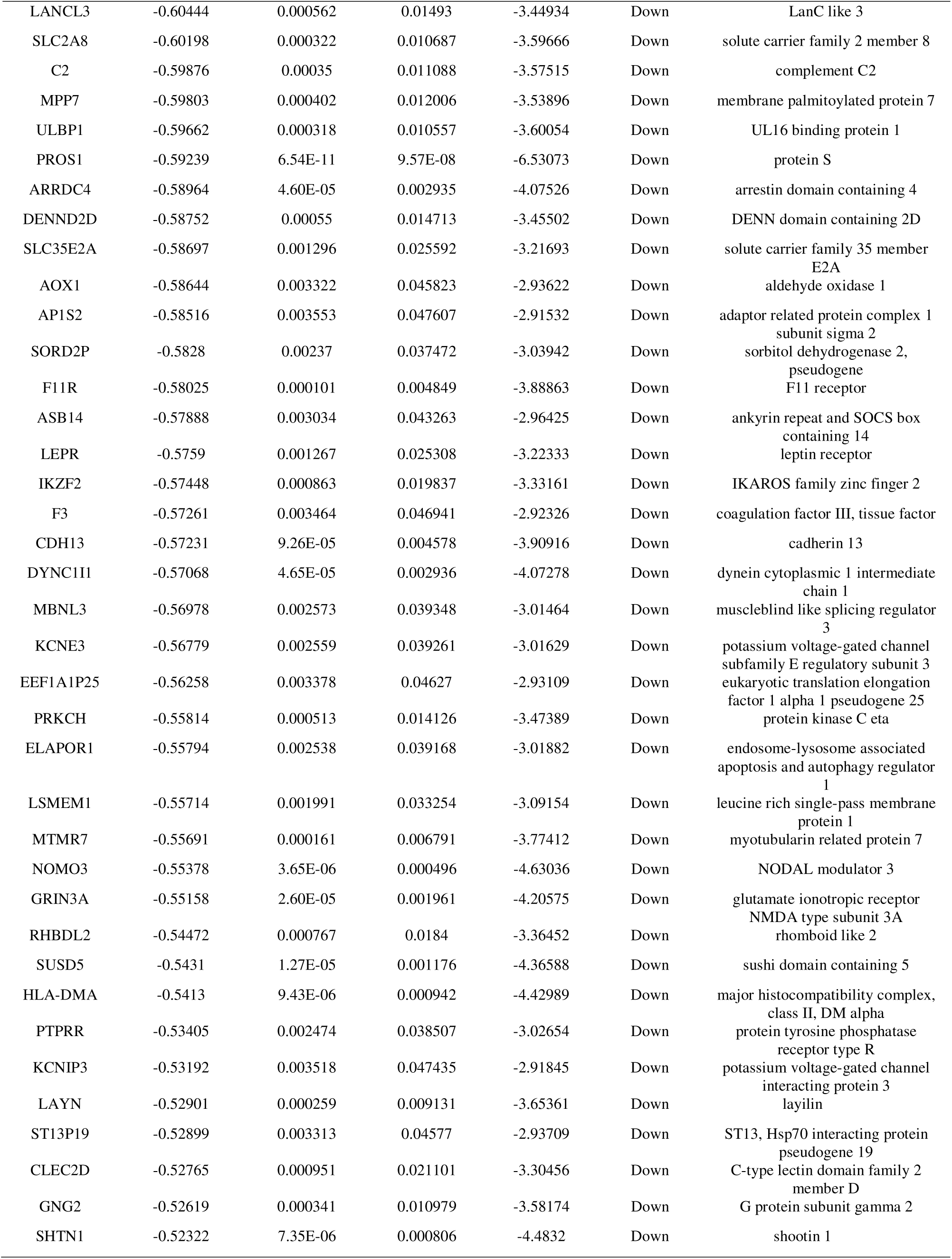
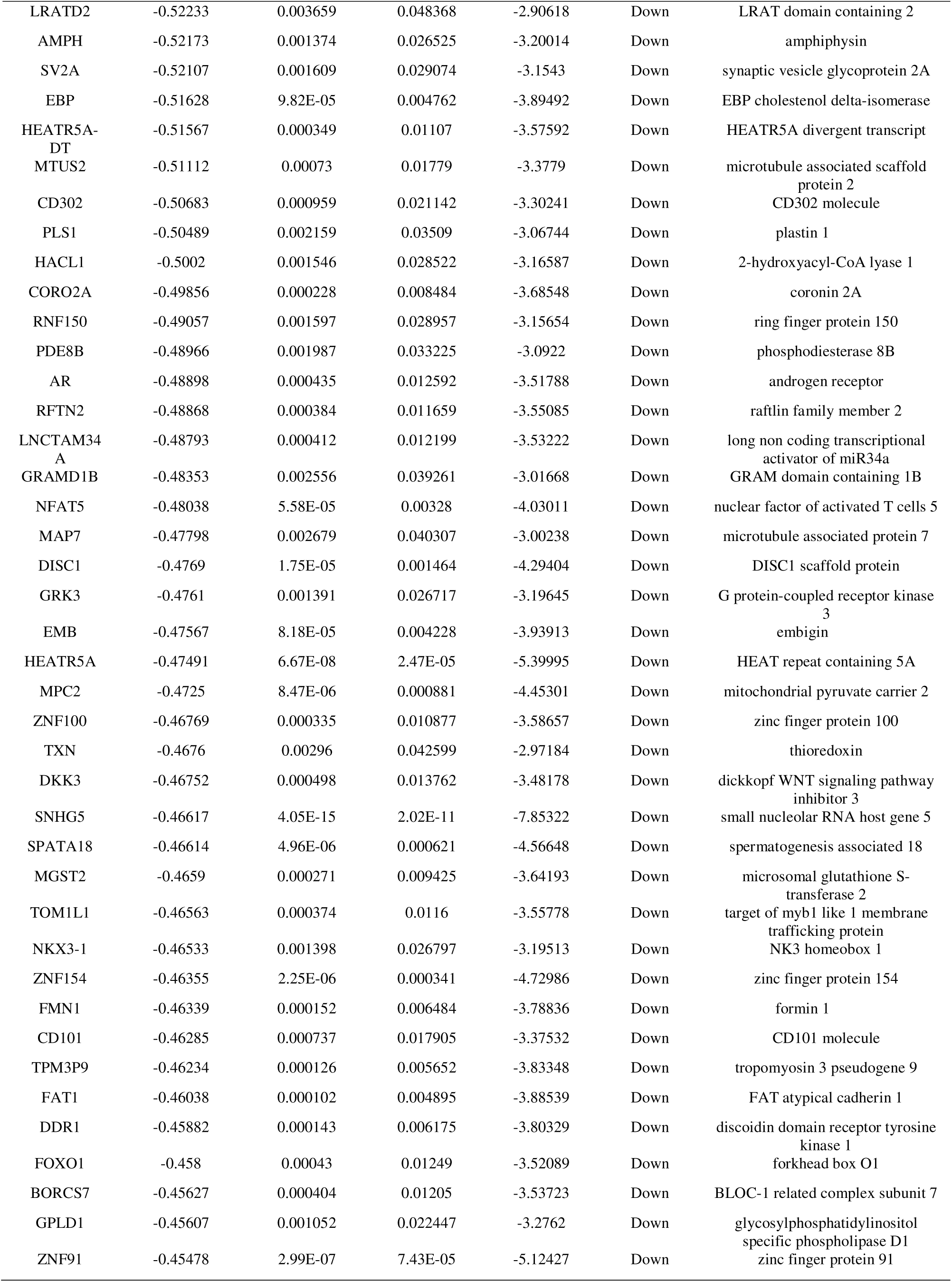
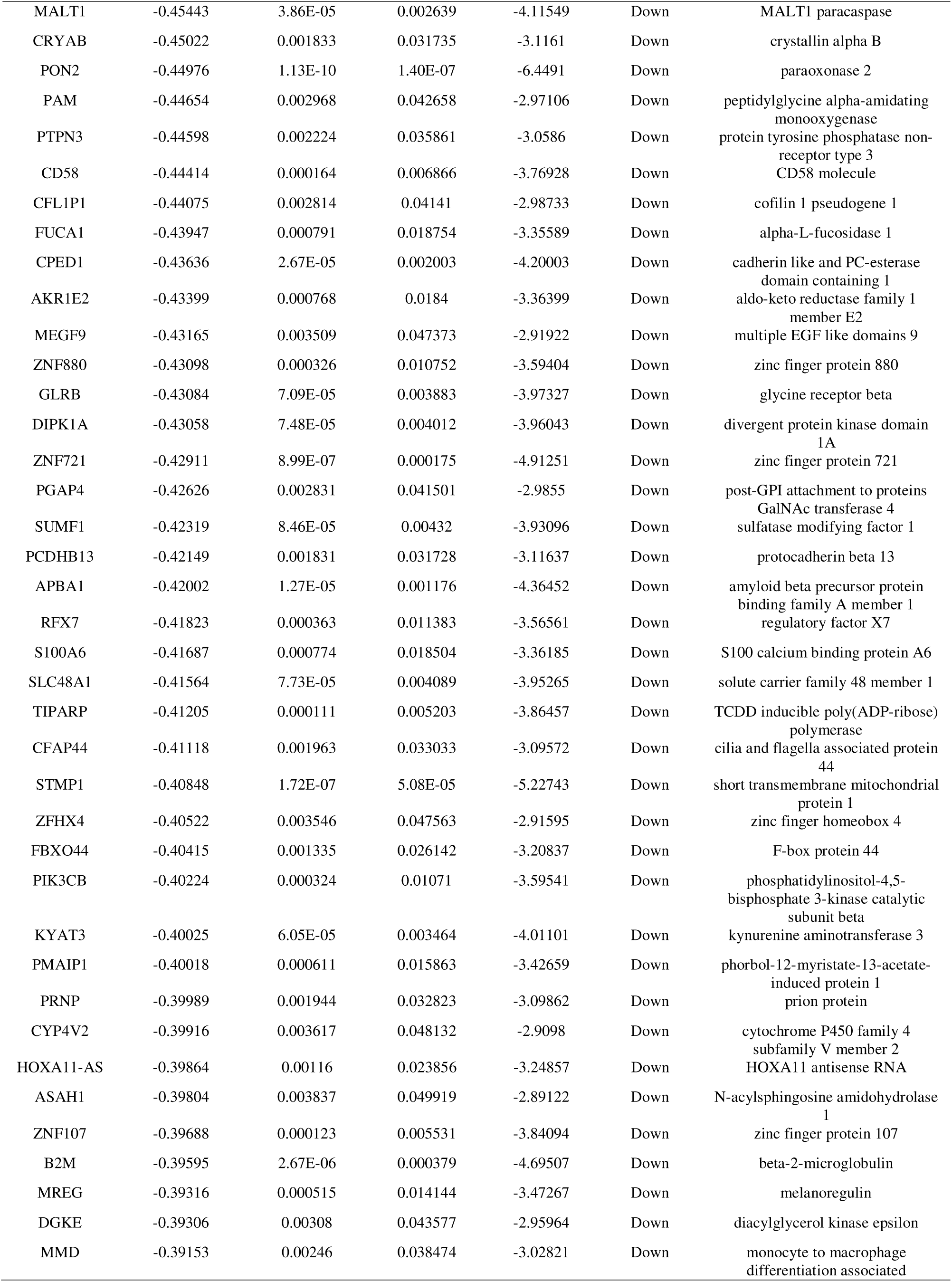
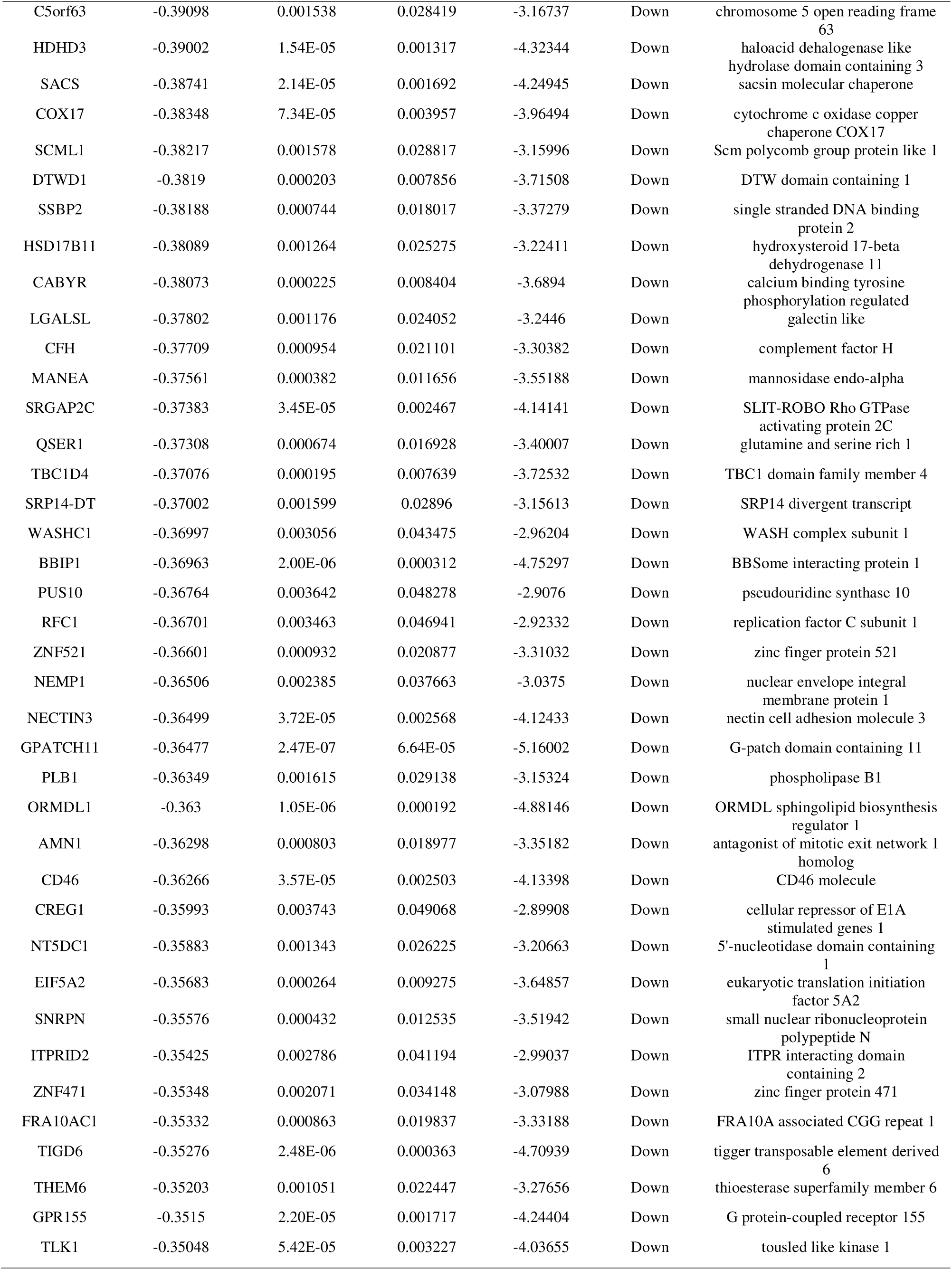
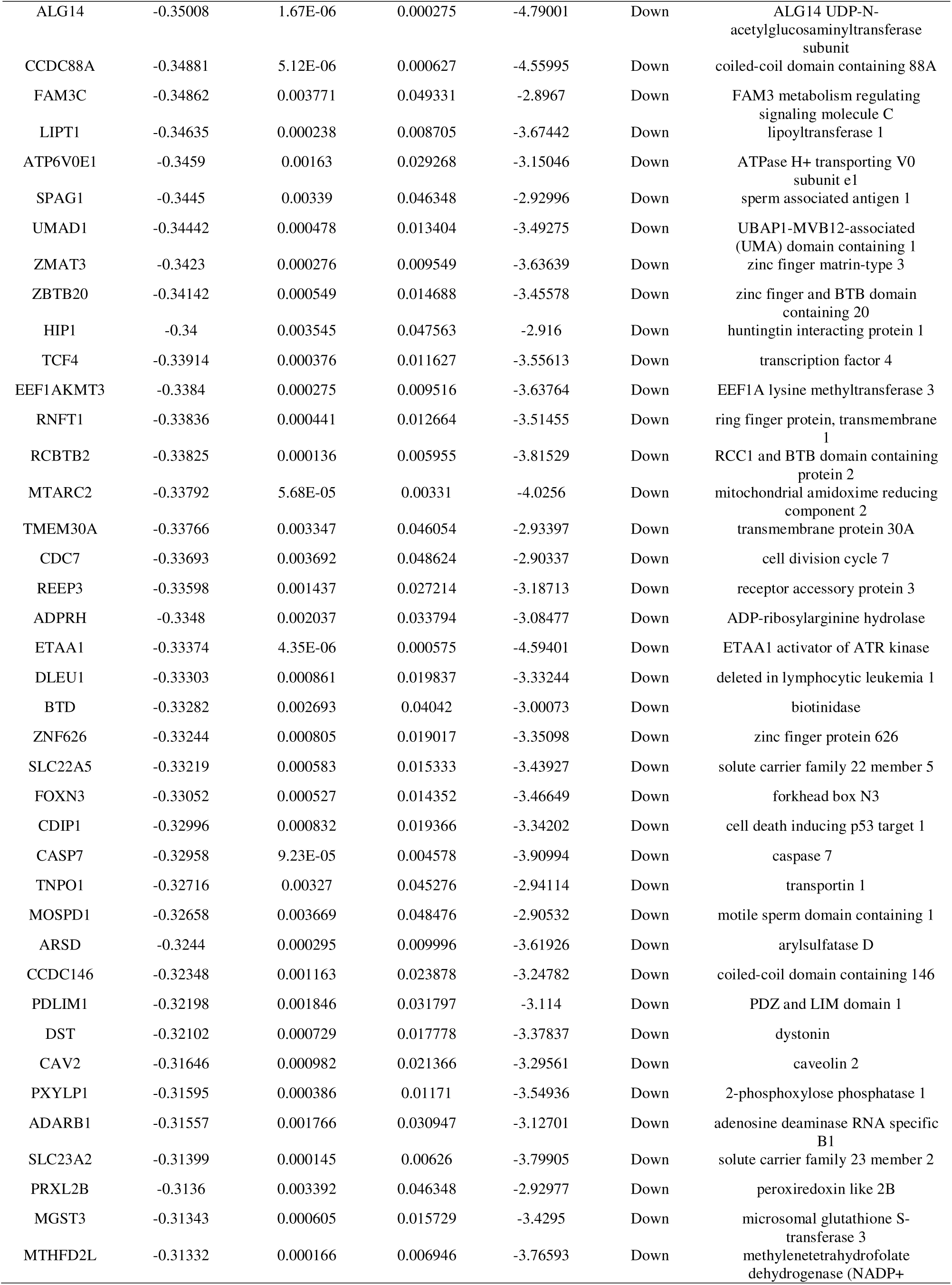
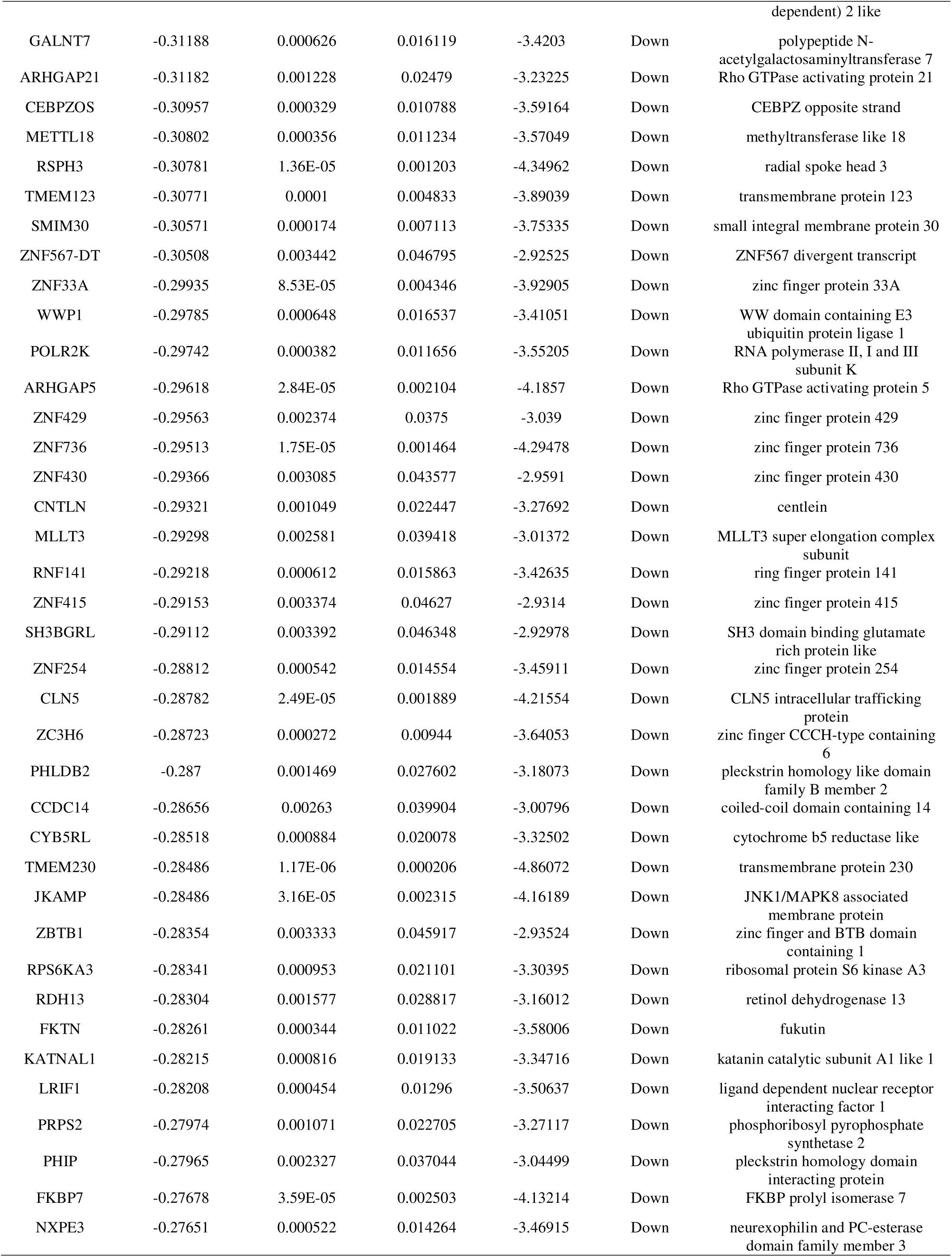
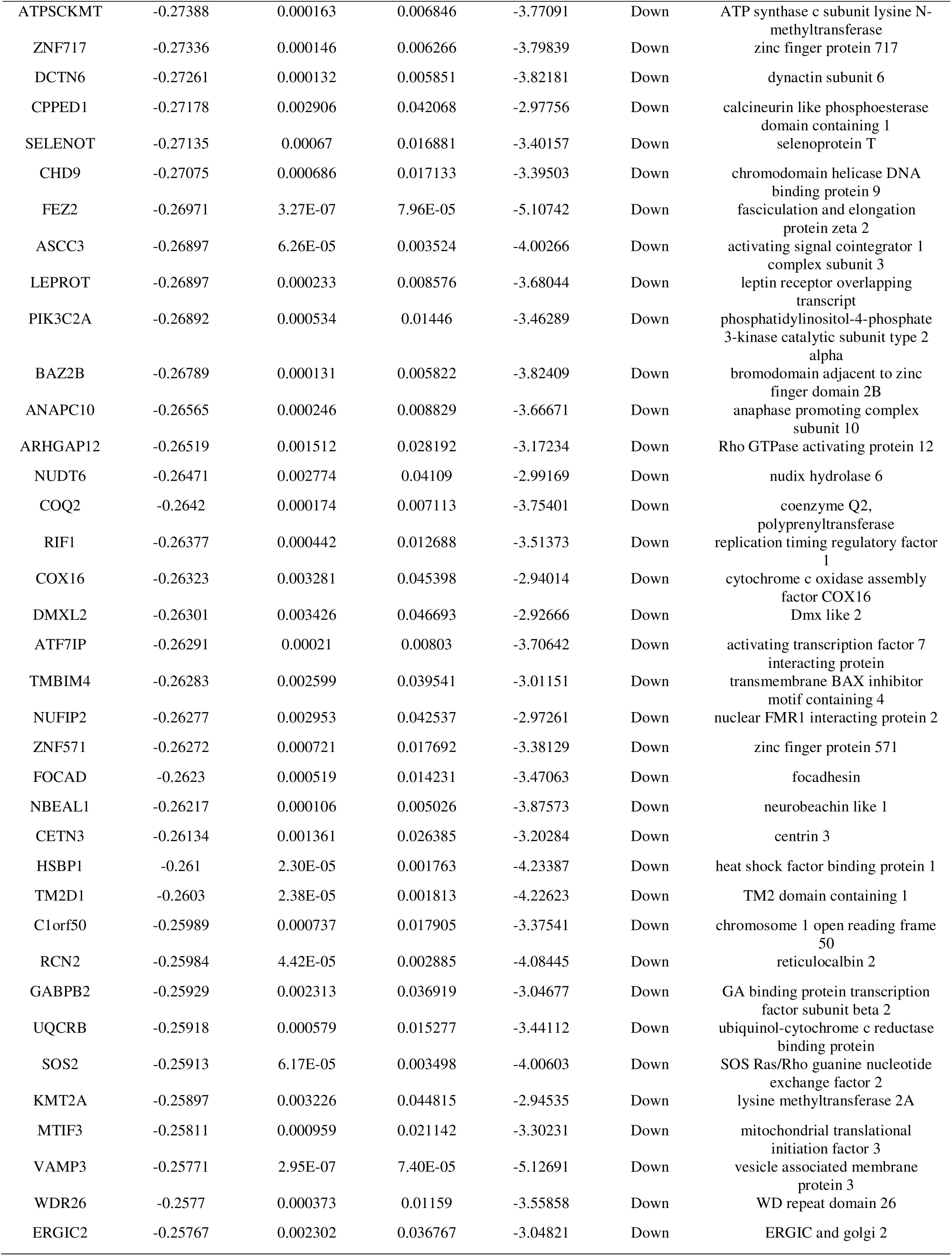
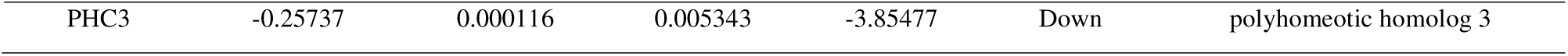
The statistical metrics for key differentially expressed genes (DEGs)

### GO and pathway enrichment analyses of DEGs

The GO ontology contains three terms: CC, MF and BP. The results demonstrated that the most significant GO terms for up regulated genes were “regulation of molecular function” (BP), “intracellular non-membrane-bounded organelle (CC) ” and “protein binding (MF)”, whereas for the down regulated genes “developmental process (BP)”, “anatomical structure development (CC)” and “cation binding (MF)” turned to be important (Table 2). Furthermore, the REACTOME pathway enrichment analysis indicated that “interferon signaling”, “cytokine signaling in immune system”, “platelet activation, signaling and aggregation”, and “hemostasis” pathways played an essential role in PCOS pathogenesis (Table 3).

**Table 2.**
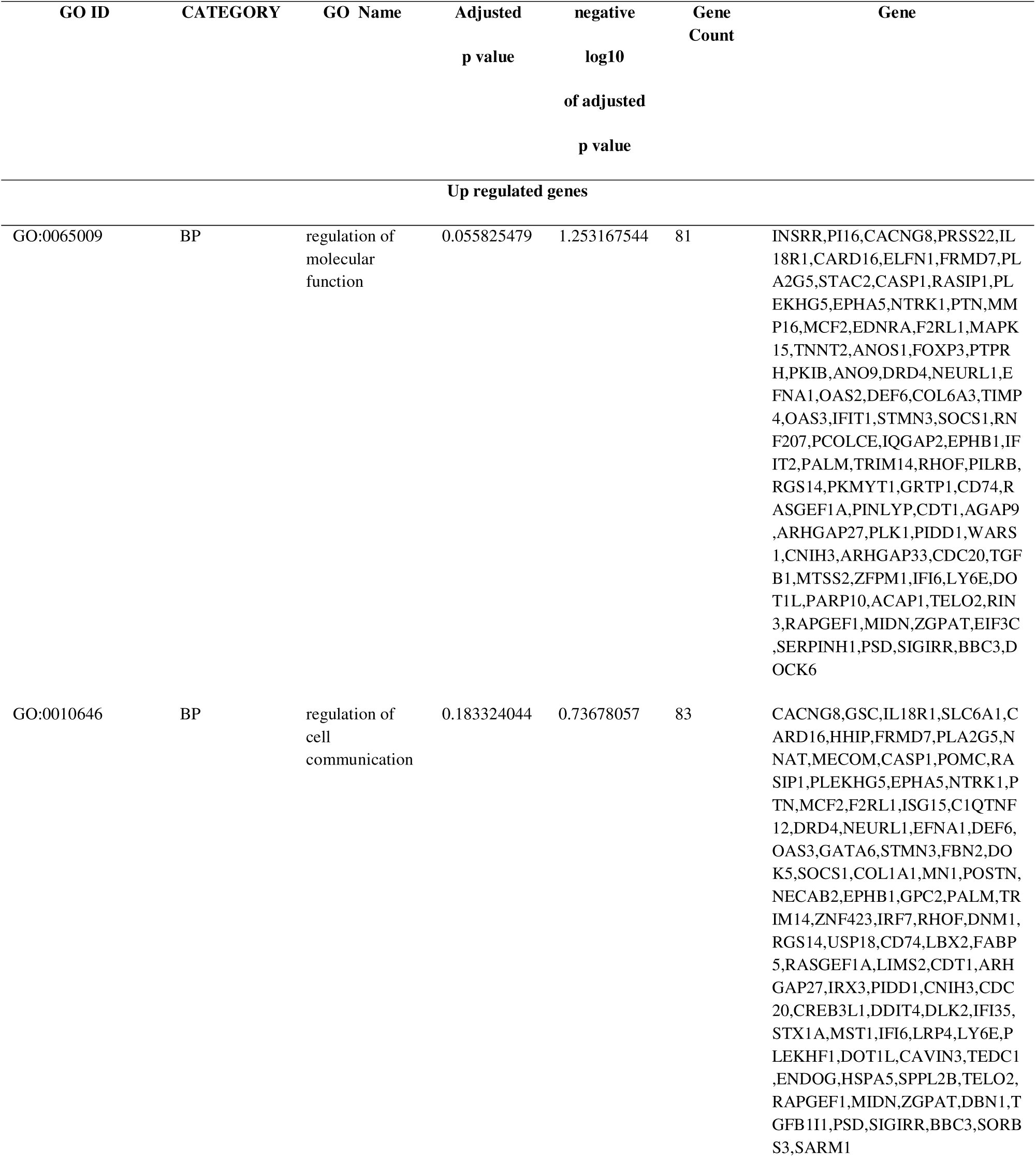
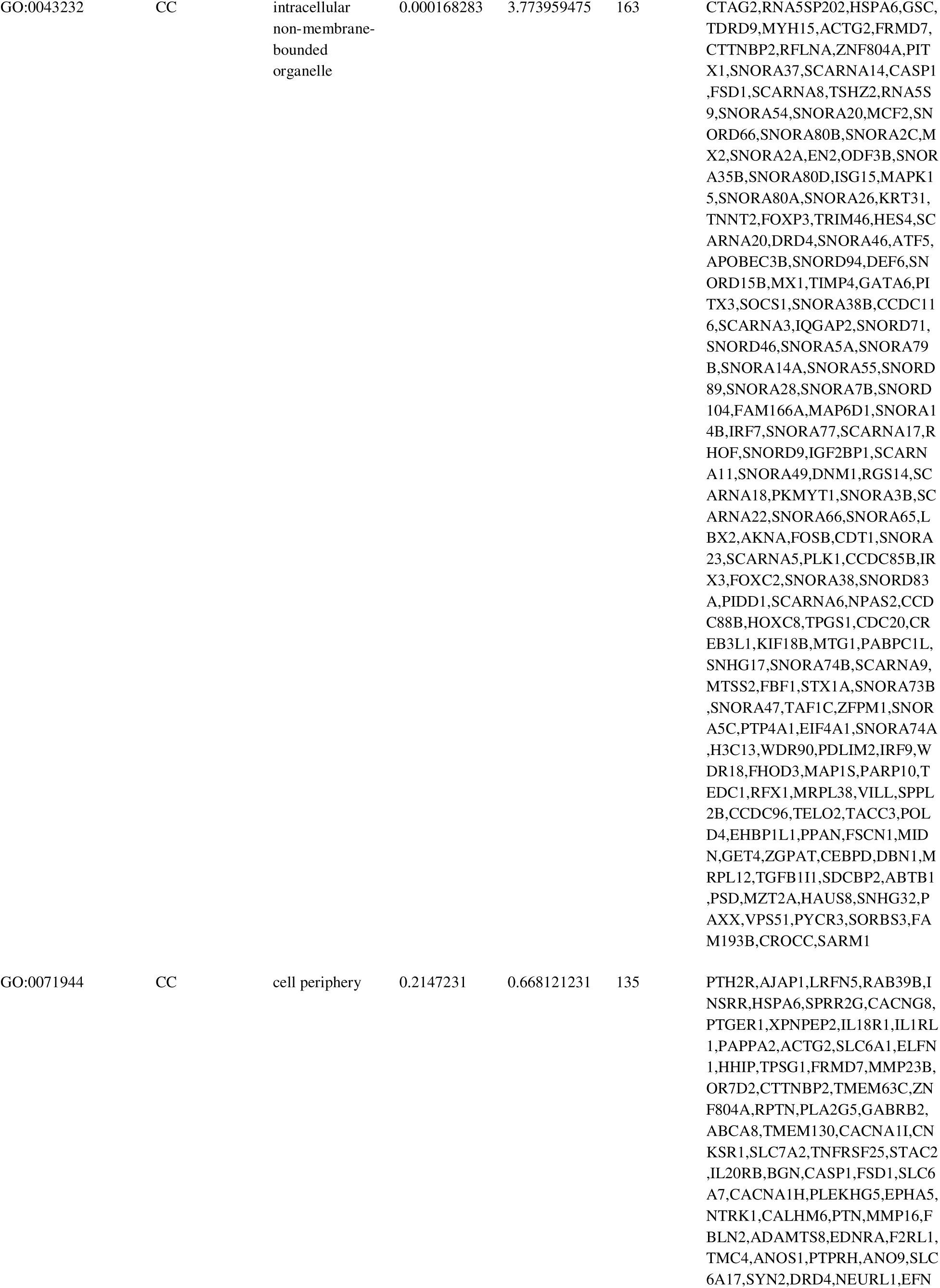
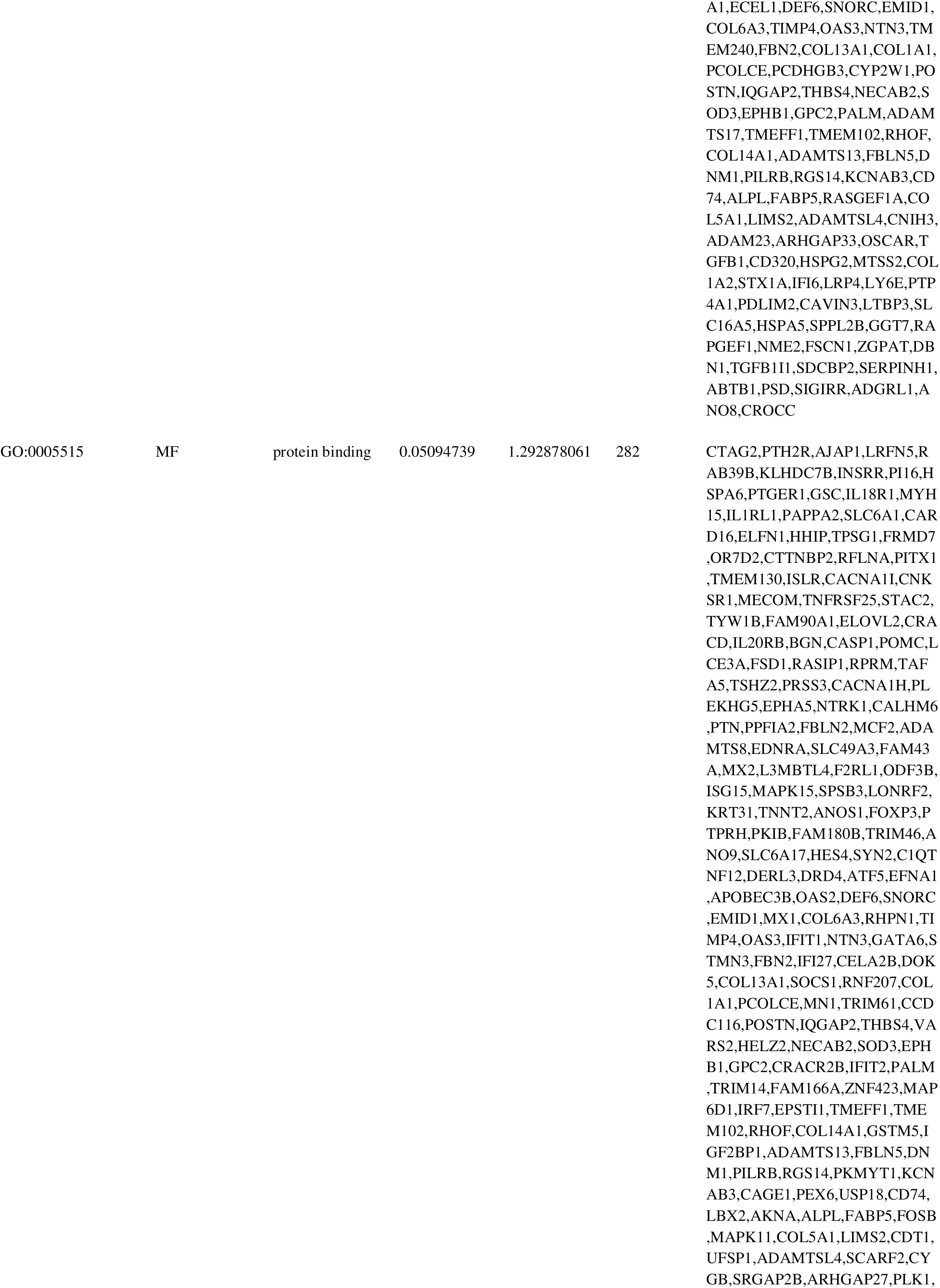
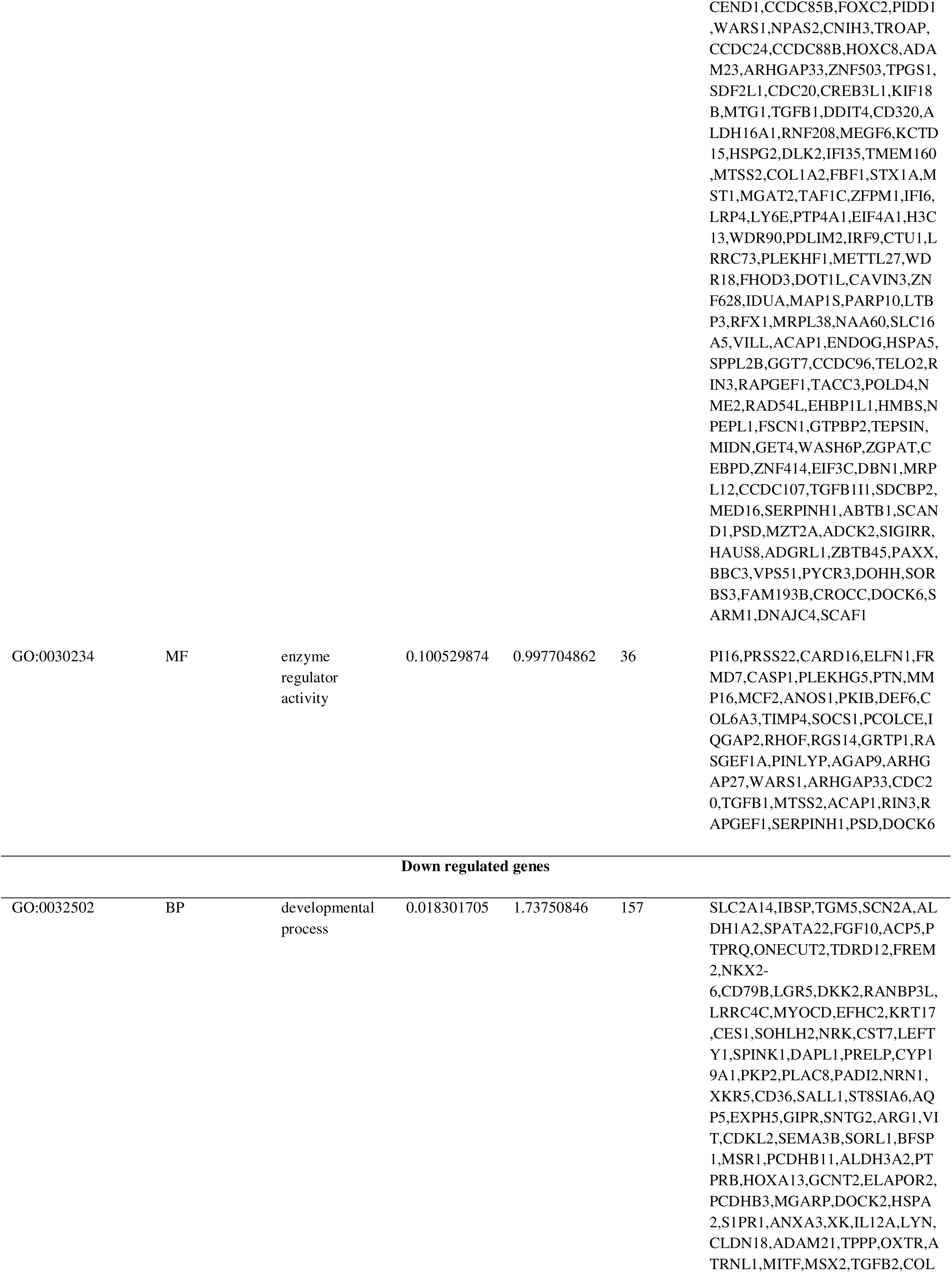
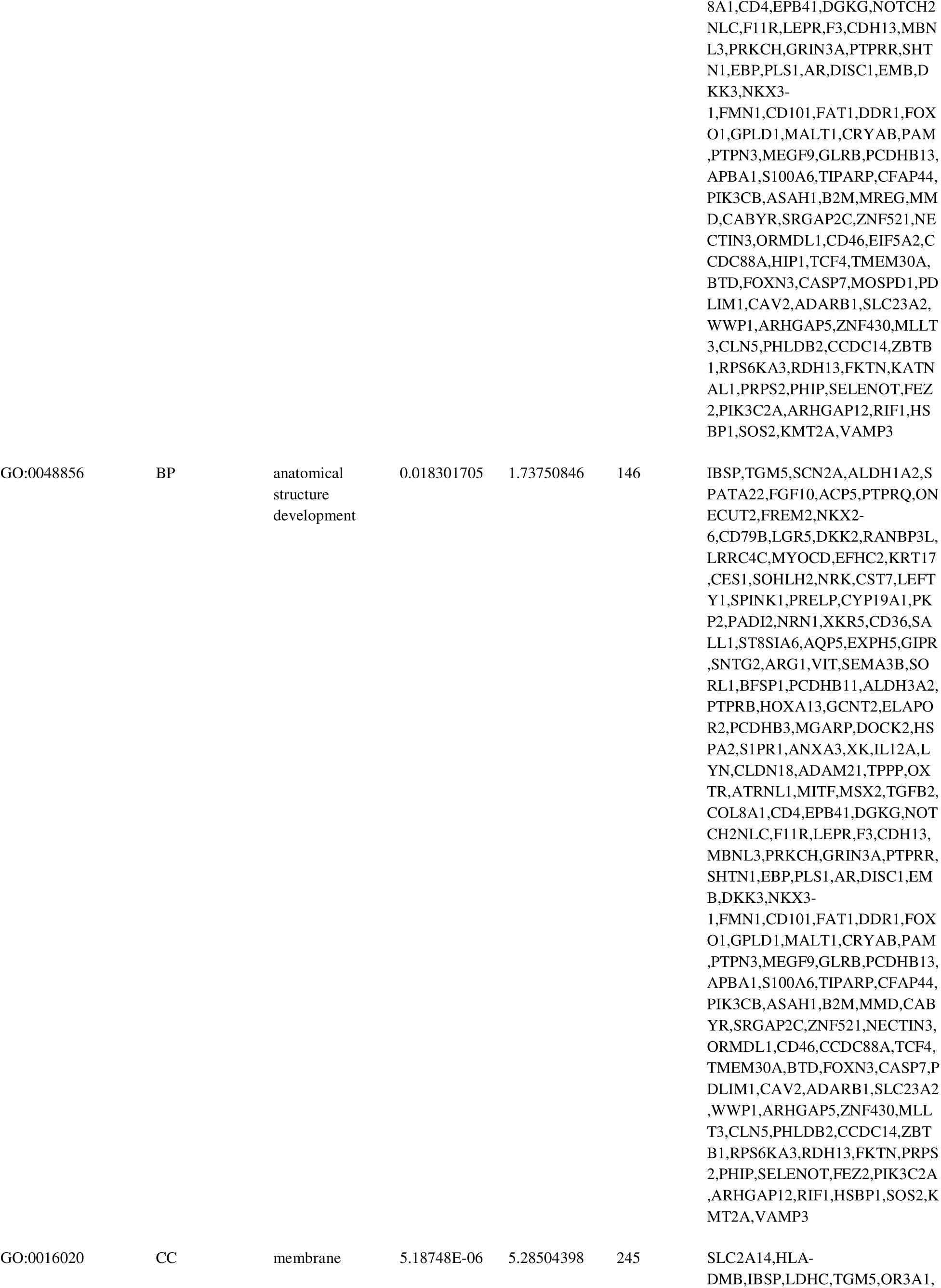
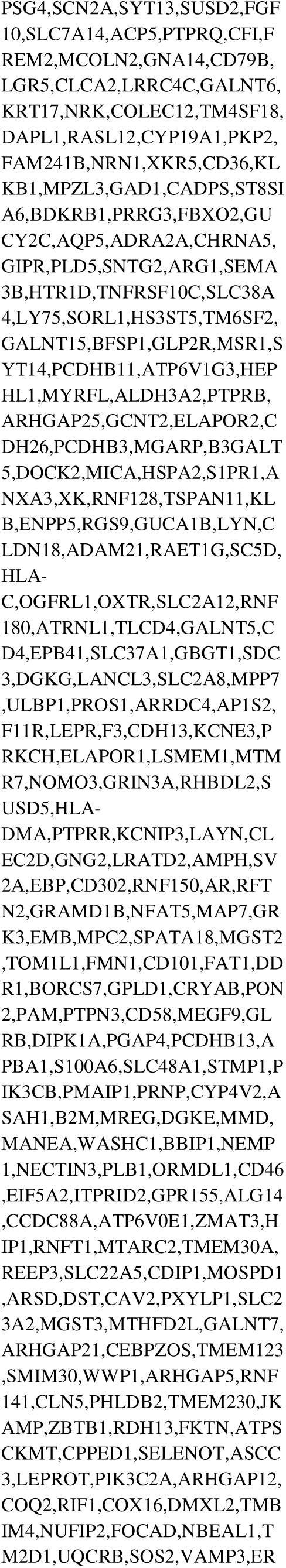
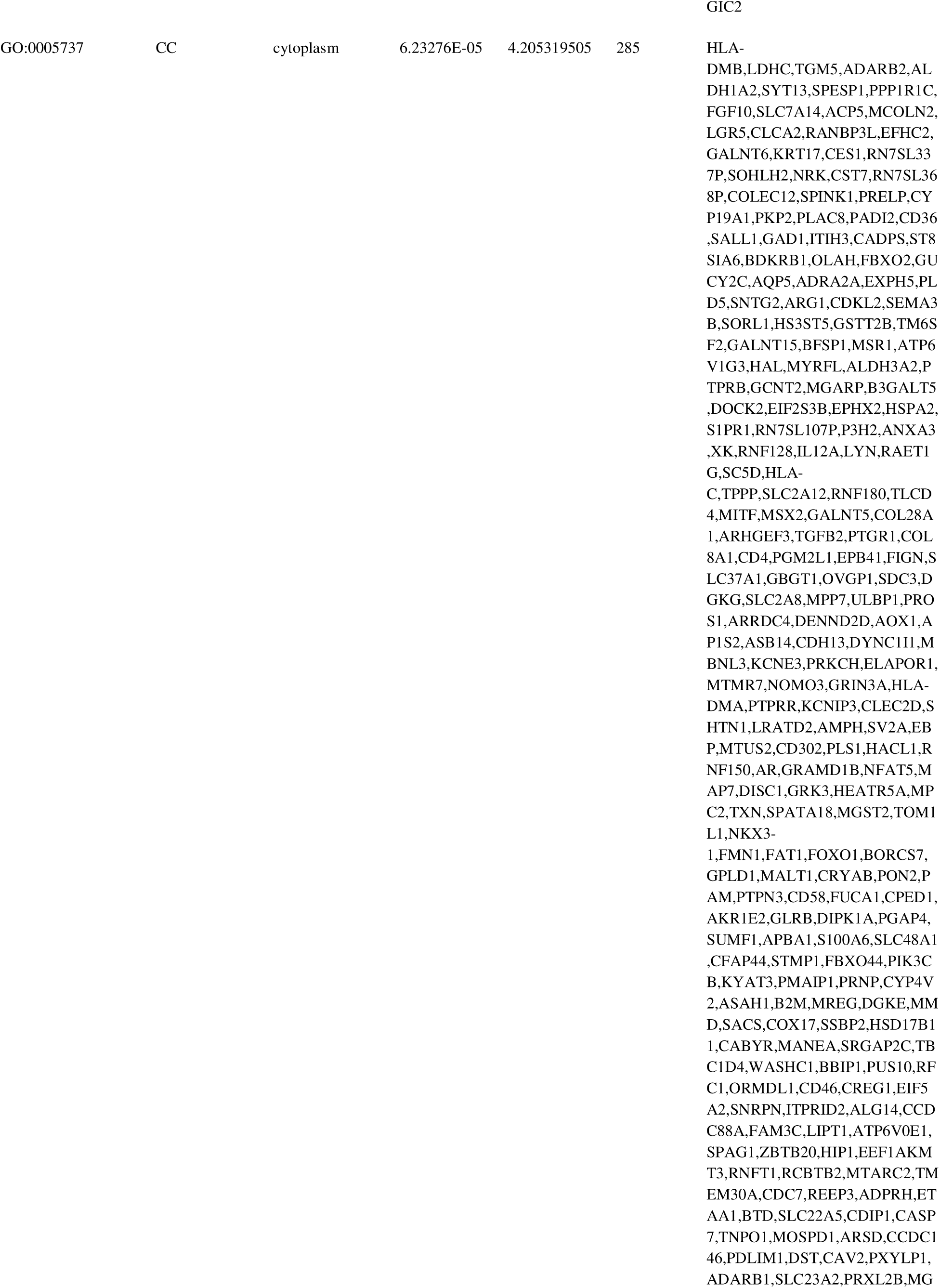
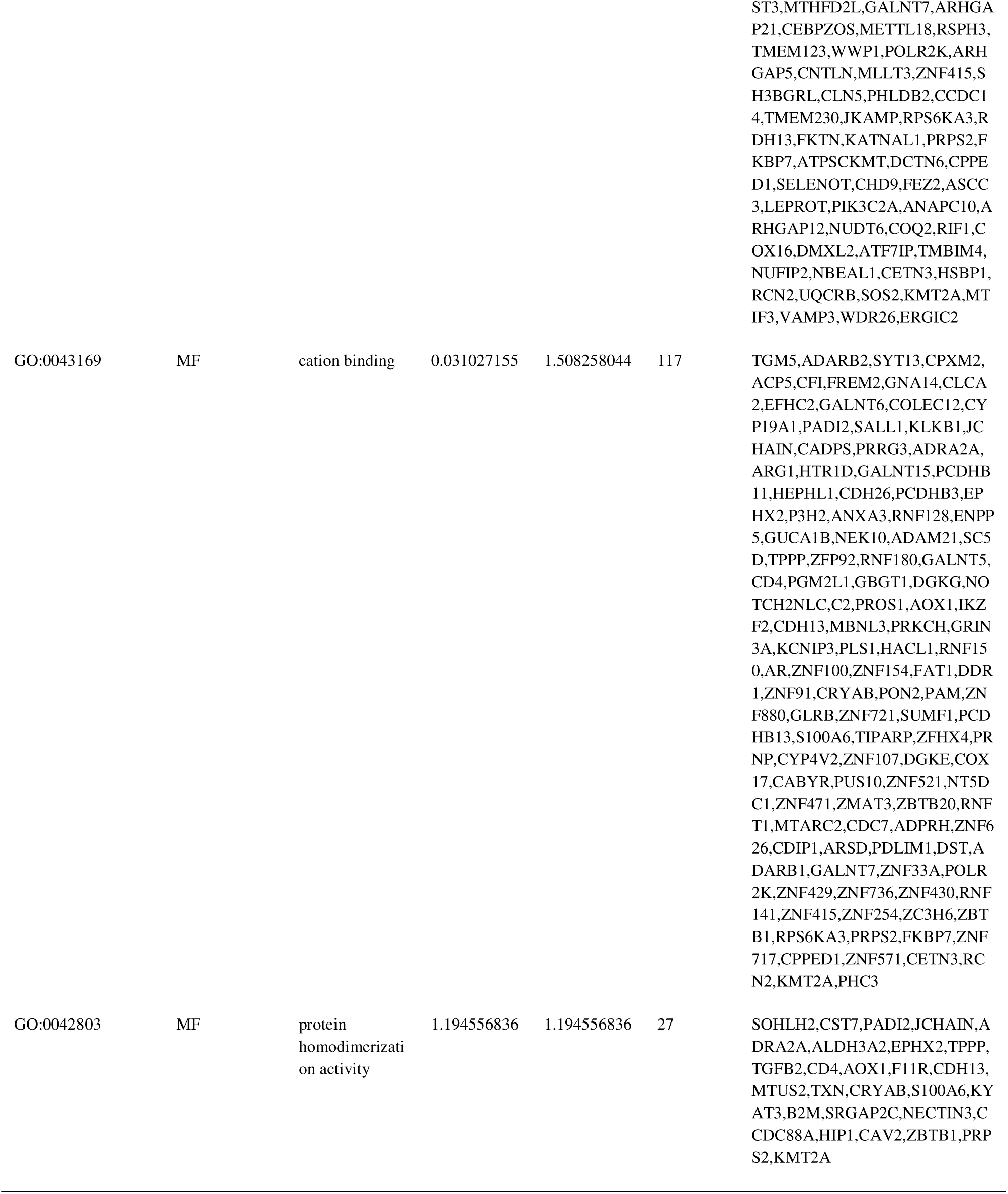
The enriched GO terms of the up and down regulated differentially expressed genes

**Table 3.**
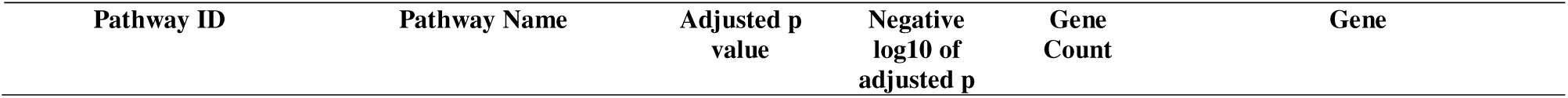
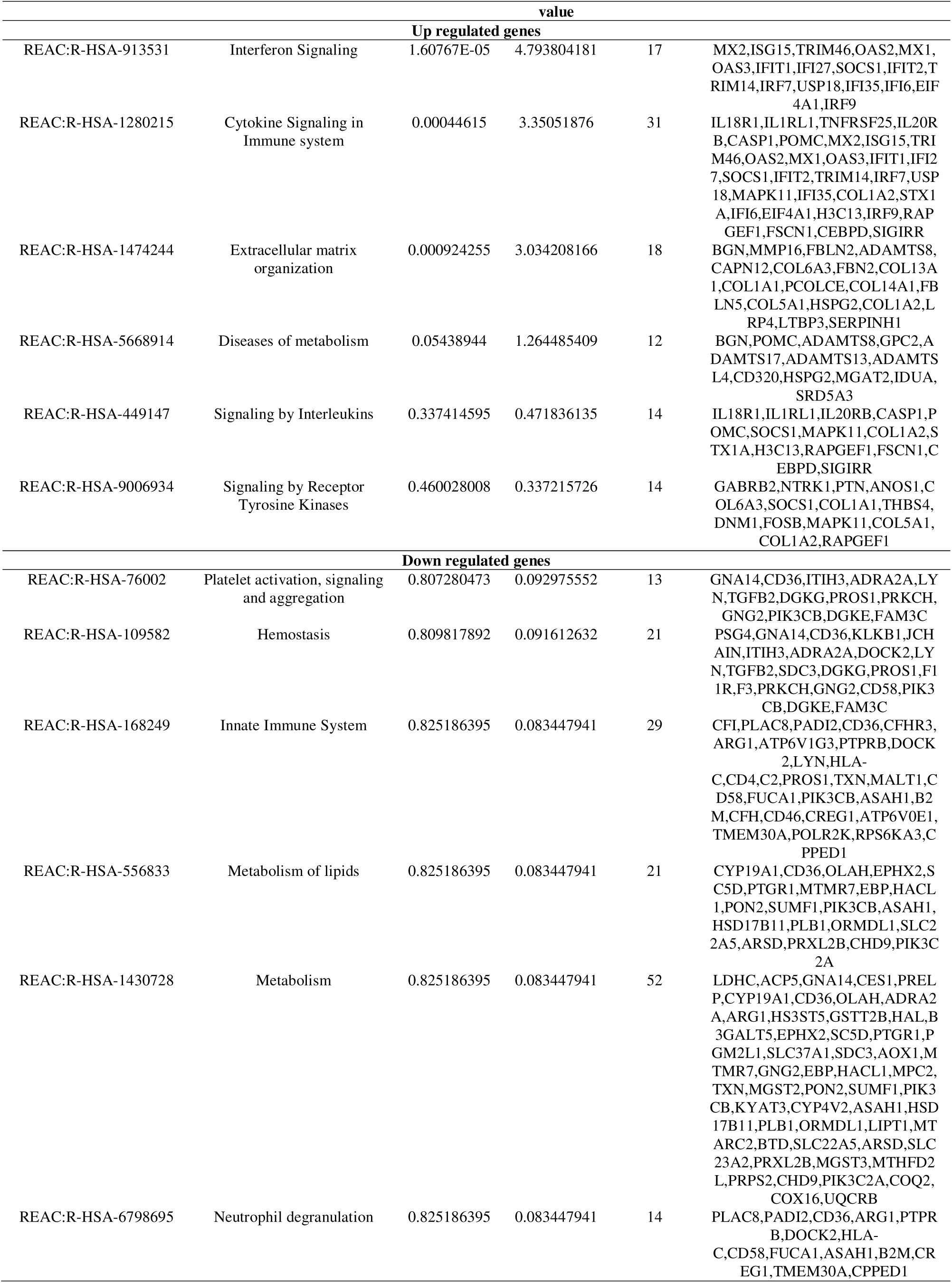
The enriched pathway terms of the up and down regulated differentially expressed genes

### Construction of the PPI network and module analysis

Via the HiPPIE interactome database website, DEGs were screened into the PPI network, which contained 5246 nodes and 8907 edges (Fig. 3) with red and blue colors representing up and down regulated genes, respectively. Hub genes therein with the highest degree, betweenness, stress and closeness score were determined using Cytoscape v. 3.9.1. The degree, betweenness, stress and closeness of hub genes is shown in Table 4. We identified hub genes, including up regulated HSPA5, PLK1, RIN3, DBN1 and CCDC85B, and down regulated DISC1, AR, MTUS2, LYN and TCF4. On the basis of the degree of importance, the 2 most significant modules were detected from the PPI network complex using the PEWCC1 plug-in, module 1 had 32 nodes and 62 edges (Fig. 4A) and module 2 had 21 nodes and 41 edges (Fig. 4B). The GO and REACTOME pathway enrichment analysis demonstrated that the module 1 primarily associated with intracellular non-membrane-bounded organelle and protein binding, whereas module 2 primarily associated with developmental process.

**Fig. 3.**
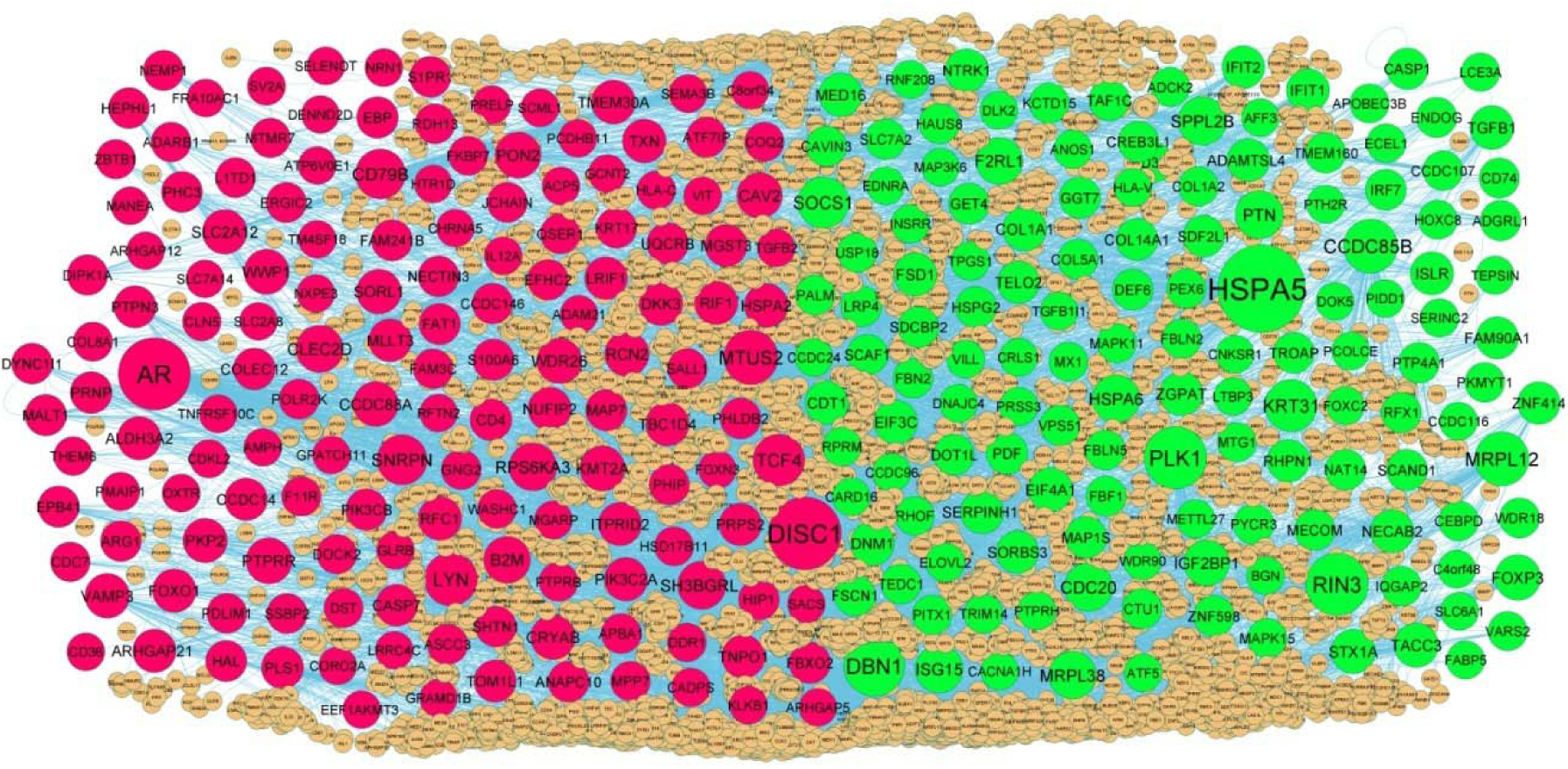
PPI network of DEGs. Up regulated genes are marked in green; down regulated genes are marked in red

**Fig. 4.**
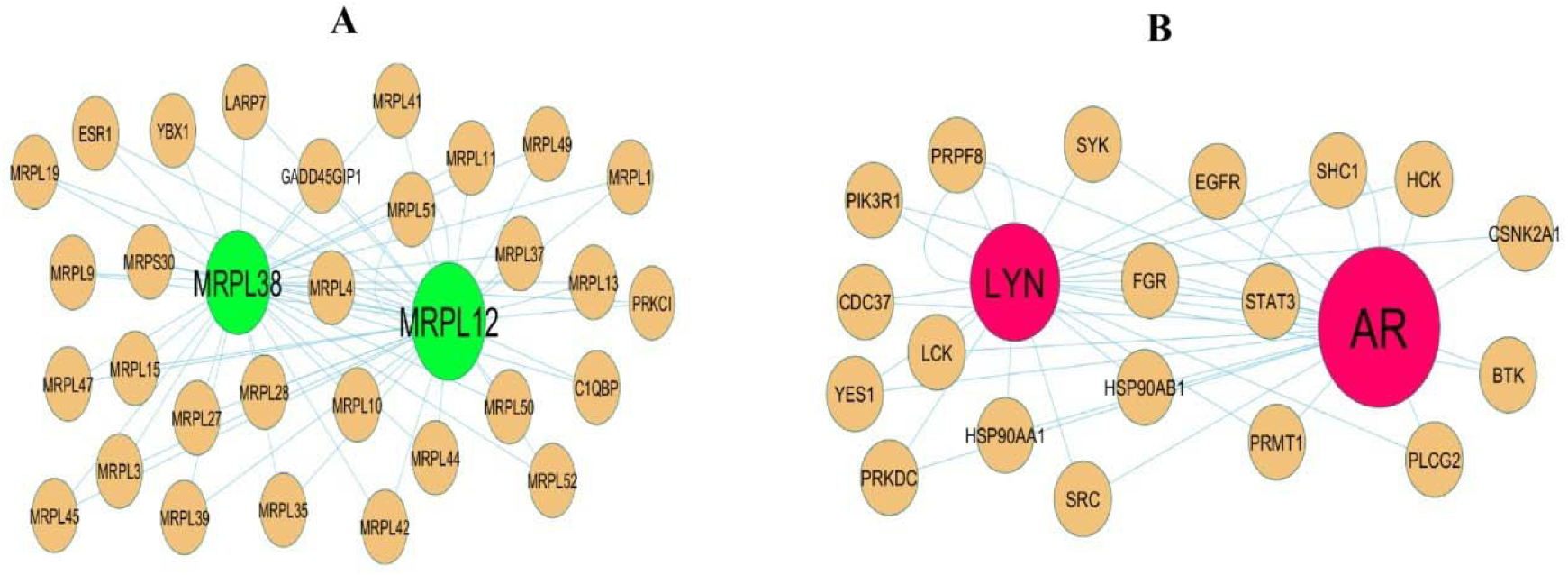
Modules selected from the DEG PPI between patients with FSGS and normal controls. (A) The most significant module was obtained from PPI network with 32 nodes and 62 edges for up regulated genes (B) The most significant module was obtained from PPI network with 21 nodes and 41 edges for down regulated genes. Up regulated genes are marked in green; down regulated genes are marked in red

**Table 4.**
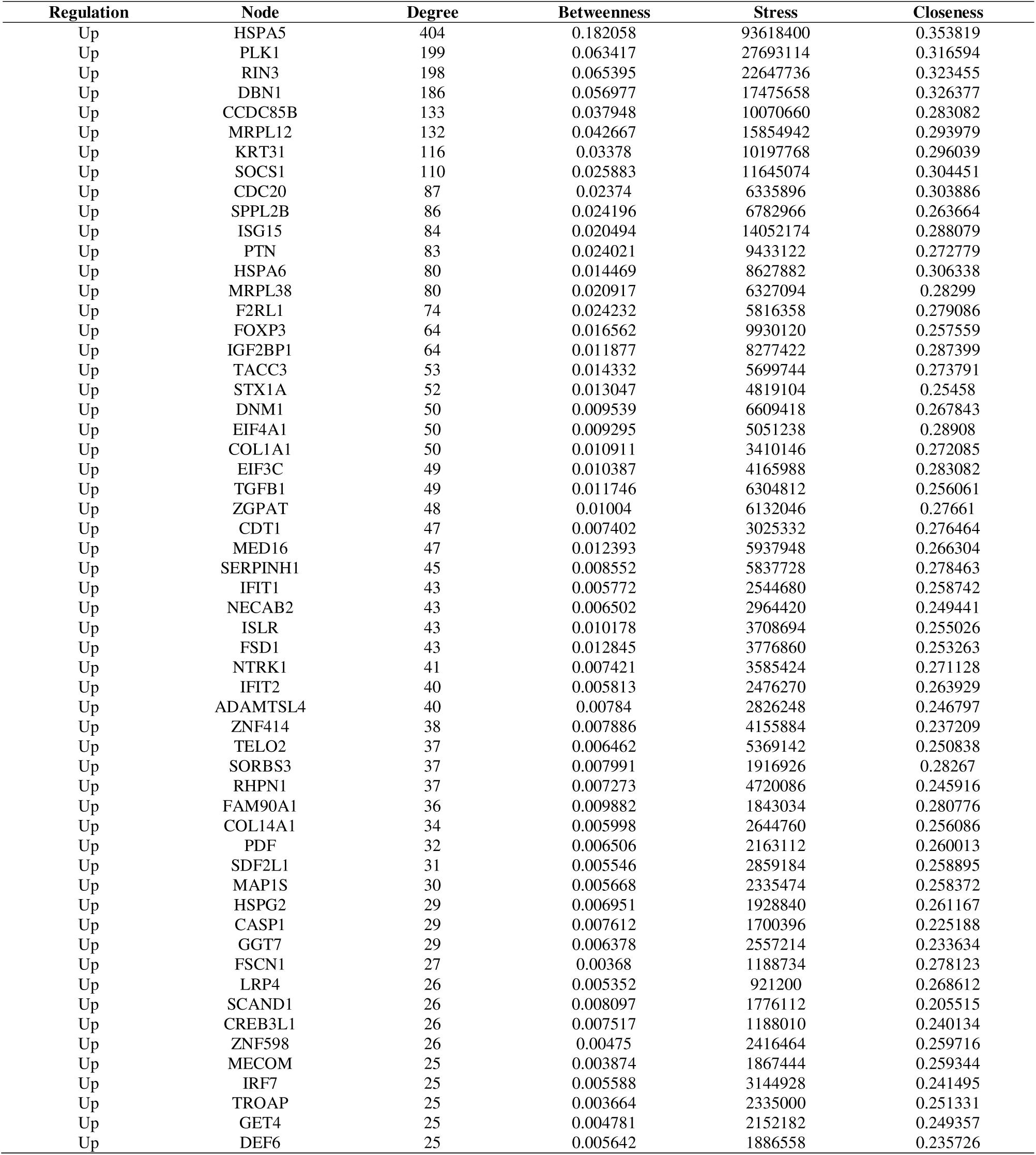
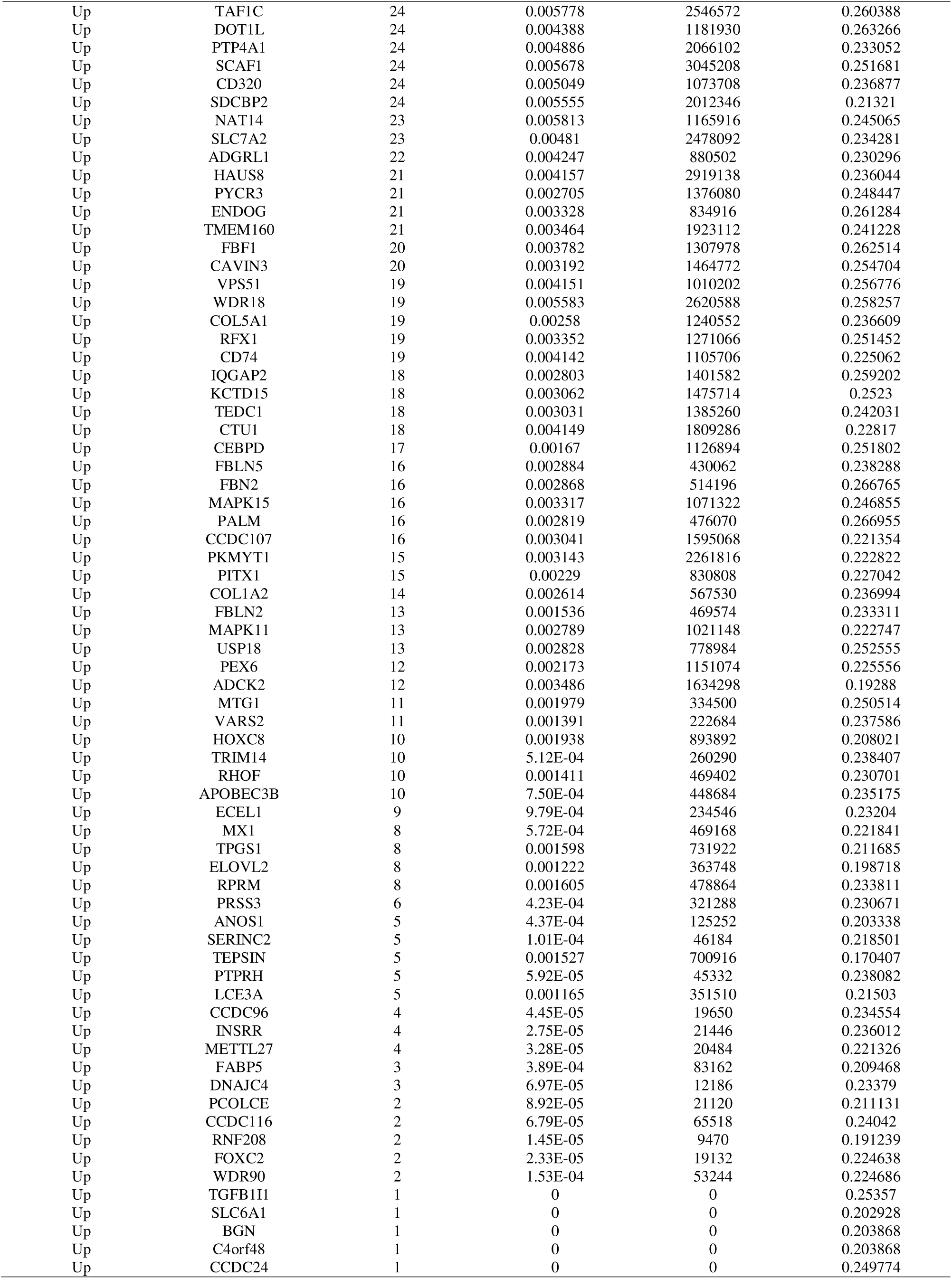
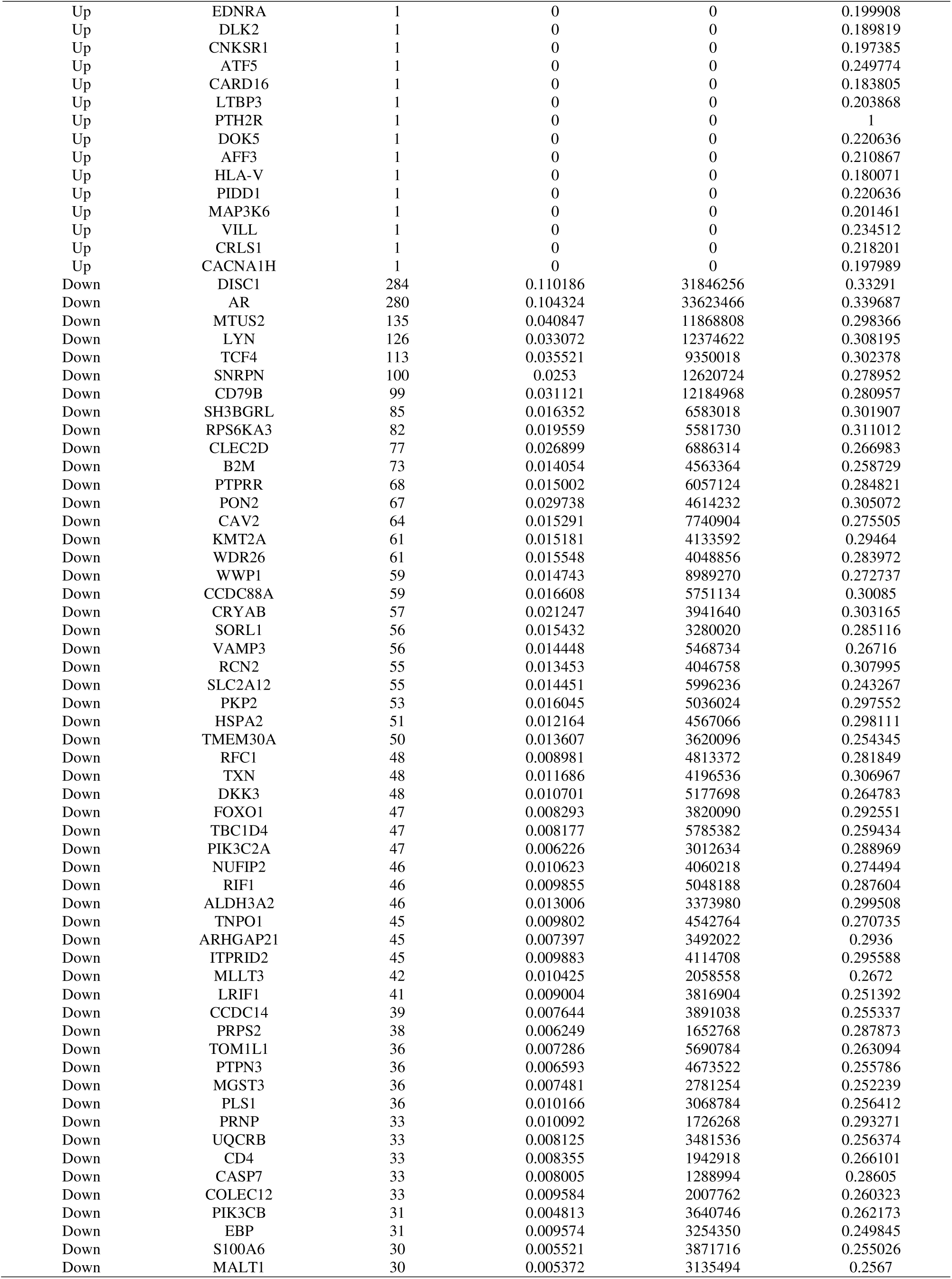
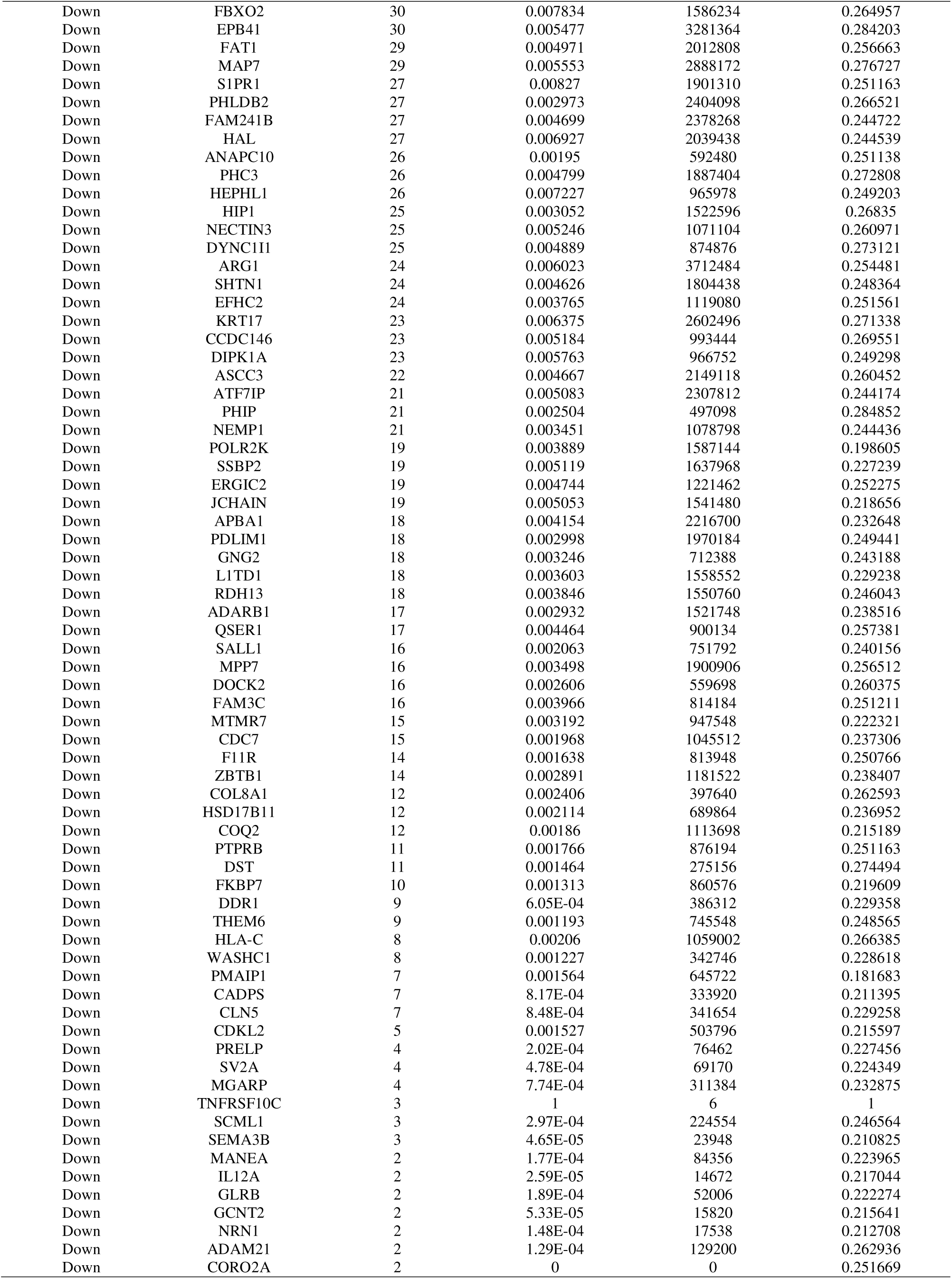
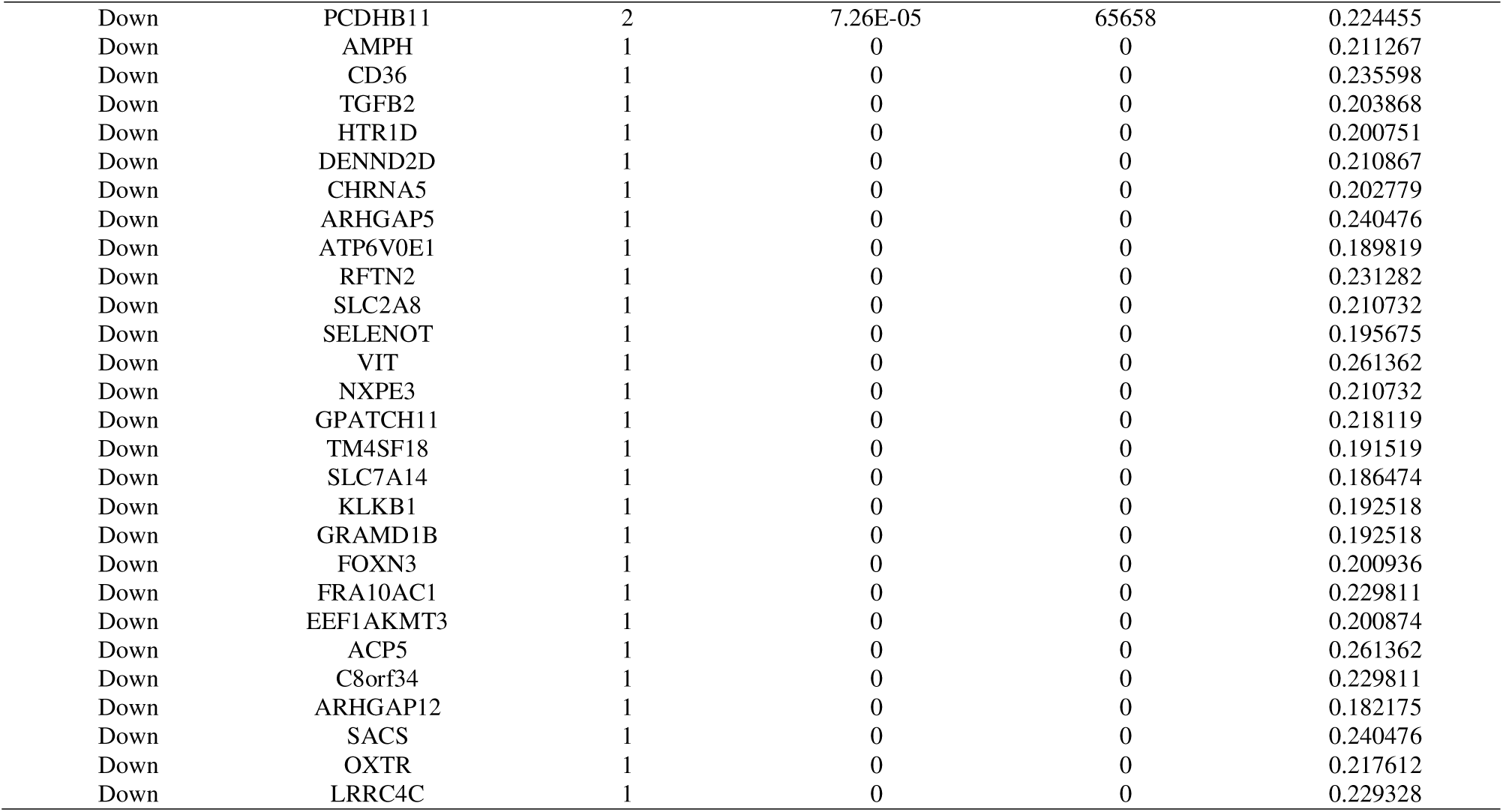
Topology table for up and down regulated genes

### miRNA-hub gene regulatory network construction

The miRNA- hub gene regulatory network miRNet database. miRNA- hub gene regulatory network was visualized, which involved 2488 nodes (286 genes and 2202 miRNAs) and 14620 edges (Fig. 5). As shown in Table 5, HSPA5 might be the targets of 123 miRNAs (ex; hsa-mir-34b-5p); MRPL12 might be the targets of 108 miRNAs (ex; hsa-mir-4728-5p); F2RL1 might be the targets of 101 miRNAs (ex; hsa-mir-302e); PLK1 might be the targets of 94 miRNAs (ex; hsa-mir-296- 5p); DBN1 might be the targets of 91 miRNAs (ex; hsa-mir-5195-5p); KMT2A might be the targets of 256 miRNAs (ex; hsa-mir-378a-5p); RPS6KA3 might be the targets of 127 miRNAs (ex; hsa-mir-5010-3p); AR might be the targets of 114 miRNAs (ex; hsa-mir-21-3p); B2M might be the targets of 107 miRNAs (ex; hsa-mir-5590-3p); TCF4 might be the targets of 87 miRNAs (ex; hsa-mir-369-3p).

**Fig. 5.**
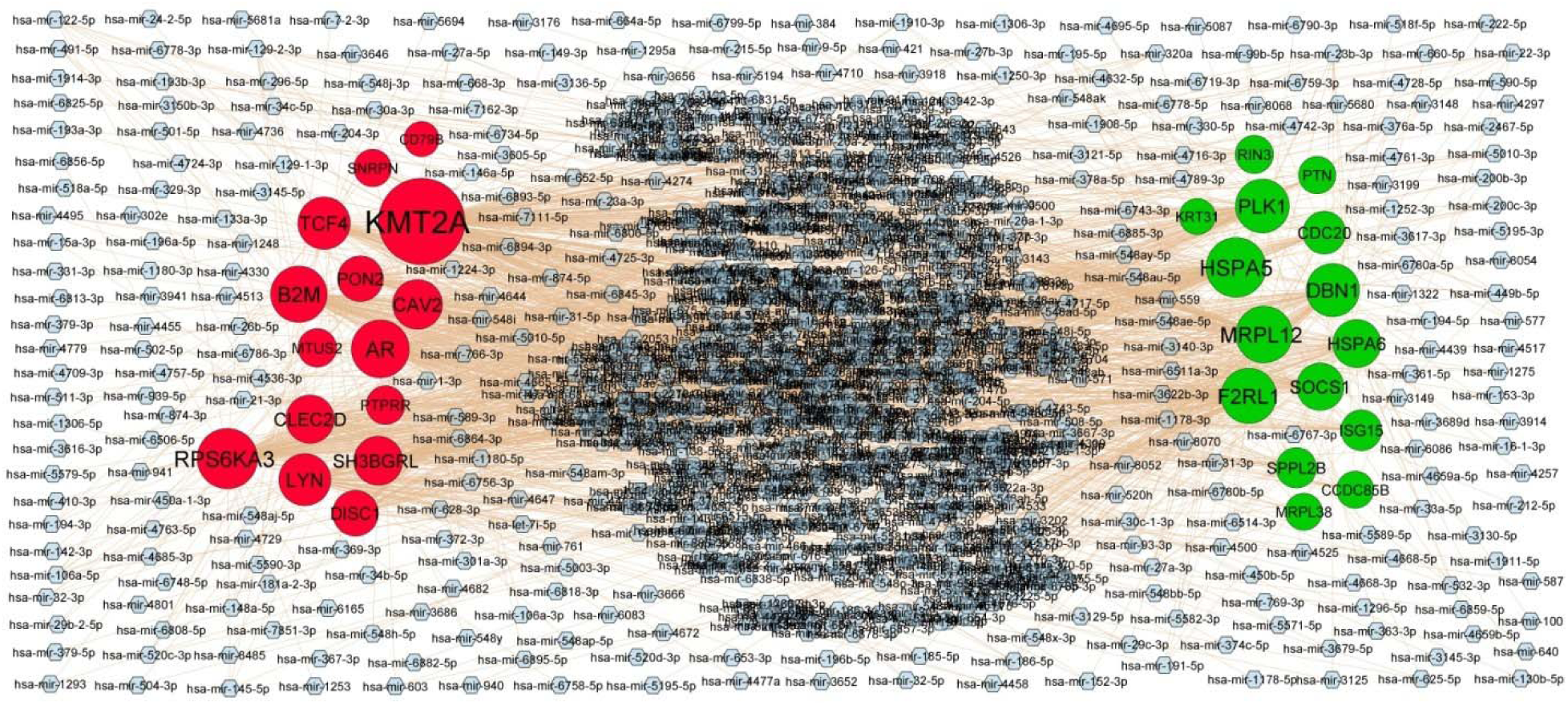
Target gene - miRNA regulatory network between target genes. The blue color diamond nodes represent the key miRNAs; up regulated genes are marked in green; down regulated genes are marked in red.

**Table 5.** miRNA - target gene and TF - target gene interaction

| Regulation | Target Genes | Degree | MicroRNA | Regulation | Target Genes | Degree | TF |
| --- | --- | --- | --- | --- | --- | --- | --- |
| Up | HSPA5 | 123 | hsa-mir-34b-5p | Up | HSPA5 | 44 | RCOR3 |
| Up | MRPL12 | 108 | hsa-mir-4728-5p | Up | RIN3 | 39 | SOX2 |
| Up | F2RL1 | 101 | hsa-mir-302e | Up | SOCS1 | 39 | GATA2 |
| Up | PLK1 | 94 | hsa-mir-296-5p | Up | MRPL12 | 36 | POU5F1 |
| Up | DBN1 | 91 | hsa-mir-5195-5p | Up | PTN | 27 | RUNX2 |
| Up | HSPA6 | 65 | hsa-mir-1178-3p | Up | F2RL1 | 27 | NANOG |
| Up | SOCS1 | 62 | hsa-mir-4779 | Up | MRPL38 | 24 | STAT3 |
| Up | CDC20 | 39 | hsa-mir-577 | Up | SPPL2B | 23 | EGR1 |
| Up | ISG15 | 29 | hsa-mir-191-5p | Up | CDC20 | 21 | CREM |
| Up | SPPL2B | 27 | hsa-mir-27a-3p | Up | PLK1 | 20 | FOXP3 |
| Up | RIN3 | 16 | hsa-mir-766-3p | Up | CCDC85B | 19 | VDR |
| Up | PTN | 12 | hsa-mir-449b-5p | Up | DBN1 | 15 | TET1 |
| Up | MRPL38 | 11 | hsa-mir-1908-5p | Up | ISG15 | 14 | IRF1 |
| Up | CCDC85B | 11 | hsa-mir-15a-3p | Up | KRT31 | 9 | AHR |
| Up | KRT31 | 5 | hsa-mir-193b-3p | Up | HSPA6 | 6 | DACH1 |
| Down | KMT2A | 256 | hsa-mir-378a-5p | Down | TCF4 | 55 | TEAD4 |
| Down | RPS6KA3 | 127 | hsa-mir-5010-3p | Down | DISC1 | 37 | DROSHA |
| Down | AR | 114 | hsa-mir-21-3p | Down | LYN | 36 | STAT5A |
| Down | B2M | 107 | hsa-mir-5590-3p | Down | AR | 33 | ESR1 |
| Down | TCF4 | 87 | hsa-mir-369-3p | Down | RPS6KA3 | 30 | PAX3 |
| Down | LYN | 85 | hsa-mir-106a-5p | Down | MTUS2 | 30 | GATA4 |
| Down | CAV2 | 74 | hsa-mir-222-5p | Down | B2M | 25 | SUZ12 |
| Down | SH3BGRL | 70 | hsa-mir-559 | Down | SNRPN | 25 | REST |
| Down | CLEC2D | 67 | hsa-mir-6818-3p | Down | PTPRR | 24 | FOXO3 |
| Down | DISC1 | 54 | hsa-mir-4458 | Down | KMT2A | 24 | MECOM |
| Down | PON2 | 53 | hsa-mir-491-5p | Down | PON2 | 24 | HNF4A |
| Down | PTPRR | 19 | hsa-mir-200c-3p | Down | CAV2 | 24 | PRDM5 |
| Down | MTUS2 | 18 | hsa-mir-4742-3p | Down | SH3BGRL | 14 | HOXB4 |
| Down | SNRPN | 13 | hsa-mir-122-5p | Down | CLEC2D | 13 | TFCP2L1 |
| Down | CD79B | 1 | hsa-mir-146a-5p | Down | CD79B | 9 | NR1H3 |

### TF-hub gene regulatory network construction

The TF- hub gene regulatory network NetworkAnalyst database. TF- hub gene regulatory network was visualized, which involved 480 nodes (293 genes and 187 TFs) and 6727 edges (Fig. 6).). As shown in Table 5, HSPA5 might be the targets of 44 TFs (ex; RCOR3); RIN3 might be the targets of 39 TFs (ex; SOX2); SOCS1 might be the targets of 39 TFs (ex; GATA2); MRPL12 might be the targets of 36 TFs (ex; POU5F1), PTN might be the targets of 27 TFs (ex; RUNX2); TCF4 might be the targets of 55 TFs (ex; TEAD4); DISC1 might be the targets of 37 TFs (ex; DROSHA); LYN might be the targets of 36 TFs (ex; STAT5A); AR might be the targets of 33 TFs (ex; ESR1); RPS6KA3 might be the targets of 30 TFs (ex; PAX3).

**Fig. 6.**
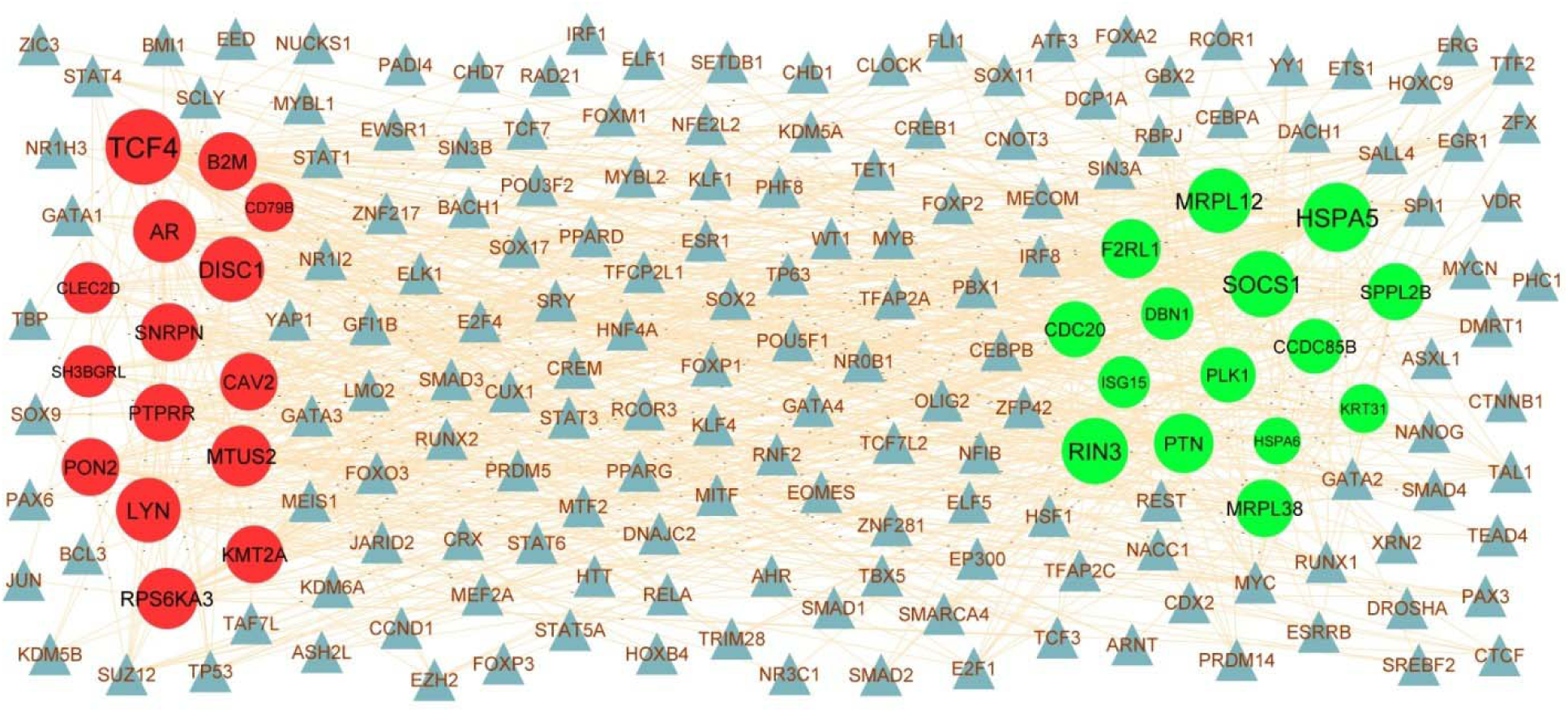
Target gene - TF regulatory network between target genes. The blue color triangle nodes represent the key TFs; up regulated genes are marked in green; down regulated genes are marked in red.

### Receiver operating characteristic curve (ROC) analysis

To determine which hub genes have the diagnose significance of PCOS patients. The ROC analyses were conducted to explore the sensitivity and specificity of hub genes for PCOS diagnosis. The ROC analysis results were available in Fig.7. Hub genes include HSPA5, PLK1, RIN3, DBN1, CCDC85B, DISC1, AR, MTUS2, LYN and TCF4 with AUC more than 0.80 were considered as hub genes, indicating that they have the capability to diagnose PCOS patients with excellent specificity and sensitivity.

**Fig. 7.**
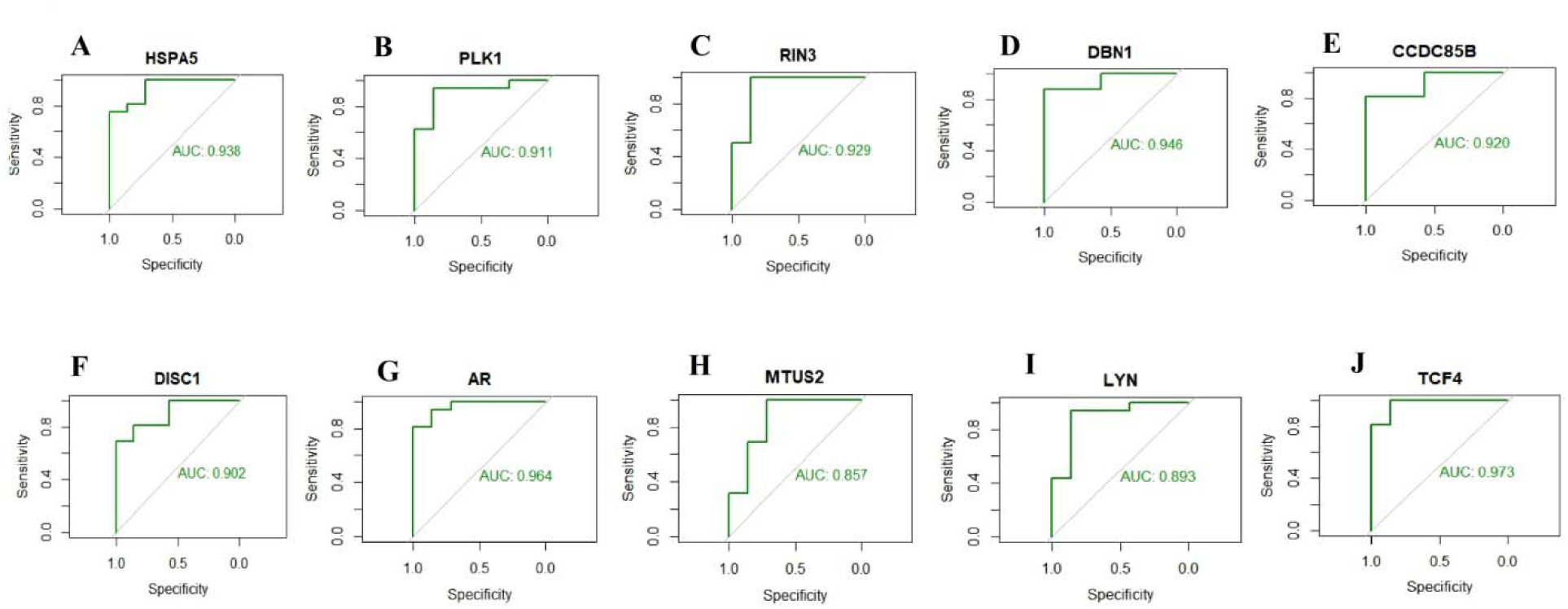
ROC curve analyses of hub genes. A) HSPA5 B) PLK1 C) RIN3 D) DBN1 E) CCDC85B F) DISC1 G) ARH) MTUS2 I) LYN J) TCF4

## Discussion

PCOS is a major global health problem due to its high prevalence and ever increasing infertility and adverse pregnancy outcomes. To discover more effective prognostic biomarkers for PCOS, a worldwide public health problem, we analyzed NGS dataset GSE199225 from the GEO database. In this study, a total of 957 DEGs were screened, including 478 up regulated genes and 479 down regulated genes. Nodin et al. [41], Xiaowei et al. [42], Xu et al. [43] and Callahan et al. [44] showed that DACH2, PTH2R, AJAP1 and HLA-DMB were associated with cancer. Johnston et al. [45] and Mangano et al. [46] reported that LRFN5 and SCN2A were important target genes of neurological and psychological problems. Deng et al. [47] found that PI16 was related to the cardiovascular problems. These data can be used to examine the DEGs that are more functionally related to PCOS development and its associated complications.

GO term and pathway enrichment analysis showed that DEGs were highly enriched in the present investigation, which undoubtedly verifies the reliability of our results. Signaling pathways include cytokine signaling in immune system [48], extracellular matrix organization [49], diseases of metabolism [50], hemostasis [51], innate immune system [52], metabolism of lipids [53] and metabolism [54] were linked with progression of PCOS. Altered expression of PLA2G5 [55], CASP1 [56], EDNRA (endothelin receptor type A) [57], F2RL1 [58], FOXP3 [59], DRD4 [60], COL6A3 [61], TIMP4 [62], SOCS1 [63], CD74 [64], TGFB1 [65], ATF5 [66], IRF7 [67], IRX3 [68], FOXC2 [69], STX1A [70], IL1RL1 [71], HHIP (hedgehog interacting protein) [72], ELOVL2 [73], BGN (biglycan) [74], POMC (proopiomelanocortin) [75], DOK5 [76], COL1A1 [77], POSTN (periostin) [78], SOD3 [79], ZNF423 [80], FABP5 [81], DDIT4 [82], KCTD15 [83], COL1A2 [84], MGAT2 [85], ENDOG (endonuclease G) [86], HSPA5 [87], CES1 [88], CYP19A1 [89], PLAC8 [90], CD36 [91], GIPR (gastric inhibitory polypeptide receptor) [92], ARG1 [93], MSR1 [94], DOCK2 [95], S1PR1 [96], OXTR (oxytocin receptor) [97], F11R [98], LEPR (leptin receptor) [99], AR (androgen receptor) [100], DKK3 [101], FOXO1 [102], PIK3CB [103], WWP1 [104], PHIP (pleckstrin homology domain interacting protein) [105], MPZL3 [106], FBXO2 [107], GUCY2C [108], ADRA2A [109], CHRNA5 [110], GLP2R [111], SDC3 [112], NFAT5 [113], PON2 [114], PRNP (prion protein) [115], DGKE (diacylglycerol kinase epsilon) [116], ARHGAP21 [117], COQ2 [118], EPHX2 [119] and FAM3C [120] were observed to be associated with the progression of obesity. Vargas-Alarcón et al [121], Frantz et al [122], Bosè et al [123], Gu et al [124], Liu et al [125], Szirák et al [126], Ye et al [127], Alikhah et al [128], Chen et al [129], Mao et al [130], Meyer et al [131], Zhang et al [132], Sun et al [133], Jiang et al [134], Yu et al [135], Jiang et al [136], Huang et al [137], Scuruchi et al [138], Wang et al [139], Ibrahim et al [140], Zha et al [141], Pan et al. [77], Dixon et al [142], Ma et al [143], Decharatchakul et al [144], Reardon et al [145], Lei et al [146], Xiang et al [147], Pan et al. [84], Hu et al. [148], Bugger and Abel [149], Ritter et al [150], Nagao et al [151], Hasson et al [152], Bampali et al [153], Roberts et al [154], Portal et al [155], Surka et al [156], Zhang et al [157], Piechota et al [158], Cannavo et al [159], Jacondino et al [160]. Tshori et al [161], Cheng et al [162], Nowzari et al [163], Choi et al [164], Ji et al [165], Huang et al [166], Zhai et al [167], Schiattarella et al [168], Brodehl et al [169], Wu et al [170], Zhao et al [171], Ujihara et al [172], Tan et al [173], Gittleman et al [174], Chen et al [175], Layrisse et al [176], Nakayama et al [177], Zhang et al [178], Vadvalkar et al [179], Burdon et al [180], Steffen et al [181], Ye et al [182], Tao et al [183] and Li et al [184] reported that PLA2G5, CASP1, TNNT2, FOXP3, DRD4, NEURL1, TIMP4, SOCS1, CD74, PLK1, ATF5, GATA6, IRF7, FOXC2, IRF9, FHOD3, BGN (biglycan), CACNA1H, FBLN2, ADAMTS8, COL1A1, POSTN (periostin), VARS2, SOD3, ADAMTS13, FABP5, ADAM23, COL1A2, MST1, ENDOG (endonuclease G), NKX2-6, MYOCD (myocardin), CES1, CYP19A1, PKP2, CD36, SALL1, ARG1, MSR1, S1PR1, OXTR (oxytocin receptor), MITF (melanocyte inducing transcription factor), MSX2, LEPR (leptin receptor), CDH13, PRKCH (protein kinase C eta), AR (androgen receptor), DKK3, FOXO1, CRYAB (crystallin alpha B), B2M, WWP1, FKTN (fukutin), PIK3C2A, KLKB1, CHRNA5, HLA-C, ARRDC4, KCNE3, MPC2, EPHX2, C2, IKZF2, ZBTB20 and RCN2 expressions might be regarded as an indicator of susceptibility to cardiovascular problems. CASP1 [185], FOXP3 [186], SOCS1 [187], GATA6 [188], IRF7 [189], POSTN (periostin) [190], CYP19A1 [191], CD36 [192], LYN (LYN proto-oncogene, Src family tyrosine kinase) [193], CD4 [194], LEPR (leptin receptor) [195], AR (androgen receptor) [196], FOXO1 [197] and PON2 [198] are a potential biomarkers for the detection and prognosis of PCOS. Udjus et al [199] Benjafield et al [200], Shetty et al. [58], Kassan et al [201], Wetzl et al [202], Le Hiress et al [203], Pal-Ghosh et al [204], Niu, [205], Fan et al [206], Deng et al [207], Chen et al [208], Sardo et al [209], Seidel et al [210], Zhu et al [211], Omura et al [212], Hu et al [213], Castoldi et al [214], Yoshida et al [215], Palao et al [216], Kušíková et al [217], Decharatchakul et al [218], Ahmed et al [219], Merklinger et al [220], Lei et al [146], Liu et al [221], Chen et al [222], Tan et al [223], Shi et al [224], Ikonnikova et al [225], Shimodaira et al [226], Pravenec et al [227], Zhang et al [228], Shah et al [229], Cicekliyurt and Dermenci [230], Ong et al. [98], Nowzari et al [163], Kim et al [231], Bonafiglia et al [232], Selle et al [233],Yoo et al [234], Kasacka et al [235], Wang et al [236], Huang et al [237], Caceres et al [238], Lei et al [239], Lu et al [240], Cui et al [241], Xiao et al [242], Hiramatsu et al [243], Oliver et al [244], Grabowski et al [245], Zhu et al [246] and Li et al [247] demonstrated that the altered expression of CASP1, EDNRA (endothelin receptor type A), F2RL1, FOXP3, TIMP4, CD74, PLK1, TGFB1, GATA6, IRF7, IRF9, BGN (biglycan), CACNA1H, FBLN2, L3MBTL4, COL1A1, POSTN (periostin), THBS4, VARS2, SOD3, ADAMTS13, FBLN5, CYGB (cytoglobin), MST1, GGT7, FGF10, CES1, CYP19A1, CD36, AQP5, ARG1, OXTR (oxytocin receptor), F11R, LEPR (leptin receptor), CDH13, DDR1, FOXO1, PAM (peptidylglycine alpha-amidating monooxygenase), S100A6, B2M, TCF4, VAMP3, GNA14, KLKB1, BDKRB1, SDC3, NFAT5, GRK3, CPXM2, EPHX2 and RCN2 are associated with hypertension. A previous study reported that CASP1 [248], EPHA5 [249], NTRK1 [250], PTN (pleiotrophin) [251], FOXP3 [252], DEF6 [253], TIMP4 [254], SOCS1 [255], CDC20 [256], TGFB1 [257], DOT1L [258], EIF3C [259], BBC3 [260], CTTNBP2 [261], PITX1 [262], EN2 [263], ISG15 [264], GATA6 [265], SNORD89 [266], IGF2BP1 [267], NPAS2 [268], SNHG17 [269], EIF4A1 [270], PDLIM2 [271], IRF9 [272], RFX1 [273], TACC3 [274], FSCN1 [275], SDCBP2 [276], SLC6A1 [277], MECOM (MDS1 and EVI1 complex locus) [278], BGN (biglycan) [279], ADAMTS8 [280], RHPN1 [281], COL1A1 [282], POSTN (periostin) [283], ZNF423 [284], TMEFF1 [285], FBLN5 [286], MAPK11 [287], COL5A1 [288], CYGB (cytoglobin) [289], TROAP (trophinin associated protein) [290], ADAM23 [291], MST1 [292], HSPA5 [293], ACP5 [294], LGR5 [295], DKK2 [296], SOHLH2 [297], SPINK1 [298], CYP19A1 [299], PADI2 [300], CD36 [301], SALL1 [302], ARG1 [303], SEMA3B [304], S1PR1 [96], CD4 [305], LEPR (leptin receptor) [306], CDH13 [307], MBNL3 [308], AR (androgen receptor) [309], DKK3 [310], NKX3-1 [311], DDR1 [312], FOXO1 [313], MEGF9 [314], S100A6 [315], PIK3CB [316], ZNF521 [317], NECTIN3 [318], CD46 [319], TCF4 [320], FOXN3 [321], ADARB1 [322], PHIP (pleckstrin homology domain interacting protein) [323], RIF1 [324], SUSD2 [325], GALNT6 [326], MPZL3 [327], FBXO2 [328], ADRA2A [329], GNG2 [330], GRAMD1B [331], NEMP1 [332], ZNF154 [333], ZFHX4 [334], ZBTB20 [335] and ZNF33A [336] are altered expression in cancer. Many studies have indicated that EPHA5 [337], NTRK1 [338], PTN (pleiotrophin) [339], MMP16 [340], EDNRA (endothelin receptor type A) [341], DRD4 [342], TIMP4 [343], EPHB1 [344], IFIT2 [345], PIDD1 [346], ARHGAP33 [347], TGFB1 [348], TELO2 [349], RIN3 [350], RAPGEF1 [351], MIDN (midnolin) [352], MYH15 [353], ZNF804A [354], ISG15 [355], ATF5 [356], DNM1 [357], NPAS2 [358], CEBPD (CCAAT enhancer binding protein delta) [359], SARM1 [360], RAB39B [361], SLC6A1 [362], ELOVL2 [363], BGN (biglycan) [364], POMC (proopiomelanocortin) [365], CACNA1H [366], FBLN2 [367], SLC6A17 [368], POSTN (periostin) [369], VARS2 [370], KCNAB3 [371], PEX6 [372], FABP5 [373], CYGB (cytoglobin) [374], CEND1 [375], HSPA5 [376], GTPBP2 [377], ADGRL1 [378], DOHH (deoxyhypusine hydroxylase) [379], ALDH1A2 [380], FGF10 [381], ACP5 [382], LRRC4C [383], EFHC2 [384], CES1 [385], CYP19A1 [386], PADI2 [387], NRN1 [388], SORL1 [389], HSPA2 [390], ANXA3 [391], IL12A [392], TPPP (tubulin polymerization promoting protein) [393], OXTR (oxytocin receptor) [394], NOTCH2NLC [395], F11R [396], LEPR (leptin receptor) [397], DISC1 [398], EMB (embigin) [399], DKK3 [400], FOXO1 [401], MALT1 [402], PAM (peptidylglycine alpha-amidating monooxygenase) [403], PIK3CB [404], ASAH1 [405], HIP1 [406], BTD (biotinidase) [407], CASP7 [408], ADARB1 [409], PHLDB2 [410], PHIP (pleckstrin homology domain interacting protein) [105], SELENOT (selenoprotein T) [411], HSBP1 [412], SOS2 [413], GAD1 [414], CADPS (calcium dependent secretion activator) [415], ADRA2A [416], SDC3 [417], AP1S2 [418], GRK3 [419], BORCS7 [420], PON2 [421], CD58 [422], PRNP (prion protein) [423], MANEA (mannosidase endo-alpha) [424], ALG14 [425], DST (dystonin) [426], ADARB2 [427], PGM2L1 [428], ZBTB20 [429], CDC7 [430] and ITIH3 [431] plays a substantial role in neurological and psychological problems. PTN (pleiotrophin) [432], FOXP3 [433], TIMP4 [434], SOCS1 [435], CDC20 [436], TGFB1 [437], GATA6 [438], SPATA22 [439], SOHLH2 [440], CYP19A1 [191], ALDH3A2 [441], HOXA13 [442], IL12A [443], CD4 [444], LEPR (leptin receptor) [195] and DMXL2 [445] were identified to be closely associated with infertility. This study indicated that enriched genes might play important role in PCOS. As previously reported, FOXP3 [446], TIMP4 [447], OAS3 [448], SOCS1 [128], CD74 [129], PLK1 [449], TGFB1 [450], RAPGEF1 [451], SERPINH1 [452], ATF5 [66], GATA6 [133], IGF2BP1 [453], IRX3 [454], FOXC2 [455], SNHG17 [456], TAF1C [457], HHIP (hedgehog interacting protein) [458], POMC (proopiomelanocortin) [459], DOK5 [76], COL1A1 [460], POSTN (periostin) [461], SOD3 [462], ADAMTS13 [463], FABP5 [464], MAPK11 [465], KCTD15 [83], HSPG2 [466], MST1 [467], MGAT2 [468], LGR5 [469], SPINK1 [470], CYP19A1 [471], PLAC8 [90], CD36 [472], AQP5 [473], GIPR (gastric inhibitory polypeptide receptor) [474], ARG1 [475], SORL1 [476], CD4 [477], F11R [478], LEPR (leptin receptor) [479], CDH13 [480], AR (androgen receptor) [481], FOXO1 [102], MALT1 [482], PAM (peptidylglycine alpha-amidating monooxygenase) [483], B2M [484], TCF4 [485], CAV2 [486], HSBP1 [487], COLEC12 [488], ADRA2A [489], SLC38A4 [490], TM6SF2 [491], GLP2R [111], KLB (klotho beta) [492], NFAT5 [493], PON2 [494], ZMAT3 [495], DST (dystonin) [426], MGST3 [496], COQ2 [118], EPHX2 [497], C2 [181] and FAM3C [120] are altered expression in type 2 diabetes mellitus. Previous studies have shown that DRD4 [498], TIMP4 [499], SOCS1 [500], CD74 [501], TGFB1 [502], ACTG2 [503], ISG15 [504], HOXC8 [505], SNHG17 [506], IL1RL1 [507], BGN (biglycan) [508], SOD3 [509], ADAMTS13 [510], CYGB (cytoglobin) [511], DDIT4 [512], CYP19A1 [513], PLAC8 [514], CD36 [515], SEMA3B [516], HOXA13 [517], OXTR (oxytocin receptor) [518], TGFB2 [519], LEPR (leptin receptor) [520], CDH13 [521], AR (androgen receptor) [522], FOXO1 [523], S100A6 [524], CD46 [525], SLC23A2 [526], GNA14 [527], HLA-C [528] and NFAT5 [529] might promote adverse pregnancy outcomes in women. COL6A3 [530], SOCS1 [531], CD74 [64], IRF7 [67], FOXC2 [69], IRF9 [532], POMC (proopiomelanocortin) [533], SOD3 [79], USP18 [534], MGAT2 [85], CD36 [535], GIPR (gastric inhibitory polypeptide receptor) [536], MSR1 [94], MSX2 [537], AR (androgen receptor) [538], FOXO1 [539], PIK3CB [103], TCF4 [540], NFAT5 [113], PON2 [541], and PRNP (prion protein) [115] were found to be involved in insulin resistance. The above investigation suggest that we can provide useful information for elucidating the development mechanism of PCOS and its complications, and searching for novel therapeutic targets and biomarkers through GO and pathway enrichment analysis.

To explore the moleculer pathogenesis of PCOS and its complications, we constructed PPI network and modules analysis for the reason that the DEGs would be grouped and ordered in the network judging by their interactions. HSPA5 [87] and AR (androgen receptor) [100] have been demonstrated to enhance obesity. Altered expression of HSPA5 [293], AR (androgen receptor) [309] and TCF4 [320] were associated with progression of cancer. Altered expression of HSPA5 [376], RIN3 [350] and DISC1 [398] were found to be substantially related to neurological and psychological problems. Altered expression of PLK1 [130] and AR (androgen receptor) [166] were easily found in cardiovascular problems. Recently, increasing evidence demonstrated that PLK1 [204] and TCF4 [237] were altered expression in hypertension. Previous studies had shown that the altered expression of PLK1 [449], AR (androgen receptor) [481] and TCF4 [485] were closely related to the occurrence of type 2 diabetes mellitus. A study had shown that regulation of AR (androgen receptor) [196] and LYN (LYN proto-oncogene, Src family tyrosine kinase) [193] promoted the PCOS. AR (androgen receptor) [522] played an important role in adverse pregnancy outcomes in women. AR (androgen receptor) [538] and TCF4 [540] have been known to be involved in insulin resistance progression. We gave a new confirmation for that DBN1, CCDC85B, MRPL38, MRPL12 and MTUS2 are expected to become novel biomarkers for PCOS prognosis. This investigation may provide reference for research the connection between PCOS and its complications.

We built a miRNA-hub gene regulatory network and TF-hub gene regulatory network of hub genes in PCOS and its complications based on the Cytoscape software. Finally, we got hub genes, miRNA and TFs of PCOS and its complications. HSPA5 [87], F2RL1 [58], AR (androgen receptor) [100], SOCS1 [63], RUNX2 [542] and ESR1 [543] were an important participant in obesity. HSPA5 [293], AR (androgen receptor) [309], TCF4 [320], SOCS1 [255], PTN (pleiotrophin) [251], hsa-mir-21-3p [544], SOX2 [545], GATA2 [546], RUNX2 [547], DROSHA [548], ESR1 [549] and PAX3 [550] have been discovered to be involved in the cancer. HSPA5 [376], RIN3 [350], PTN (pleiotrophin) [339], DISC1 [398], hsa-mir-378a-5p [551], hsa-mir-5010-3p [552] and hsa-mir-21-3p [553] are believed to be associated with neurological and psychological problems. F2RL1 [58], B2M [236], TCF4 [237], hsa-mir-34b-5p [554], hsa-mir-21-3p [455], SOX2 [456], GATA2 [557], RUNX2 [558], TEAD4 [559], DROSHA [560], STAT5A [561] and ESR1 [562] are believed to be associated with hypertension. A previous study found that increased PLK1 [130], B2M [170], SOCS1 [128], AR (androgen receptor) [166], hsa-mir-21-3p [563], GATA2 [564], RUNX2 [565] and ESR1 [566] expression were associated with cardiovascular problems. PLK1 [449], AR (androgen receptor) [481], B2M [484], TCF4 [485], SOCS1 [128], hsa-mir-21-3p [567], SOX2 [568], RUNX2 [569] and ESR1 [570] molecular markers plays important regulatory roles in type 2 diabetes mellitus. The expression of the AR (androgen receptor) [196], SOCS1 [187], LYN (LYN proto-oncogene, Src family tyrosine kinase) [193], POU5F1 [571], RUNX2 [572], DROSHA [573] and ESR1 [574] molecular markers are correlated with PCOS. AR (androgen receptor) [538], TCF4 [540], SOCS1 [531] and ESR1 [575] were found to be involved in insulin resistance. SOCS1 [435], PTN (pleiotrophin) [432], SOX2 [576] and ESR1 [577] have been demonstrated to be involved in infertility. AR (androgen receptor) [522], SOCS1 [500], hsa-mir-21-3p [578], GATA2 [579], TEAD4 [580], DROSHA [581] and ESR1 [582] contributes to the progression of adverse pregnancy outcomes in women. Our findings suggested MRPL12, DBN1, KMT2A, RPS6KA3, hsa-mir-4728-5p, hsa-mir-302e, hsa-mir-296-5p, hsa-mir- 5195-5p, hsa-mir-5590-3p, hsa-mir-369-3p and RCOR3 as novel diagnostic biomarkers for PCOS and its complications.

In summary, we identified genes differentially expressed in PCOS compared with normal control and explored their potential function and relevant pathways in the pathogenesis of PCOS and its complications. We also identify that hub genes are linked with PCOS and its complications. Meanwhile, hub genes and key pathways are independent prognostic factors for PCOS and its complications. We expect that subsequent studies will confirm the hub genes identified here as essential biomarkers for PCOS and its complications.

## Acknowledgement

I thank Ali Altintas, Copenhagen University, NNF Center for Basic Metabolic Research, Integrative Physiology, Blegdamsvej 3B, Copenhagen, Denmark, very much, the author who deposited their NGS dataset GSE199225, into the public GEO database.

## Ethical approval

This article does not contain any studies with human participants or animals performed by any of the authors.

## Informed consent

No informed consent because this study does not contain human or animals participants.

## Availability of data and materials

The datasets supporting the conclusions of this article are available in the GEO (Gene Expression Omnibus) (https://www.ncbi.nlm.nih.gov/geo/) repository. [(GSE199225) https://www.ncbi.nlm.nih.gov/geo/query/acc.cgi?acc=GSE199225]

## Consent for publication

Not applicable.

## Author Contributions

B. V. - Writing original draft, and review and editing

C. V. - Software and investigation

